# Enucleation of the *C. elegans* embryo revealed the mechanism of dynein-dependent spacing between microtubule asters

**DOI:** 10.1101/2023.07.21.549990

**Authors:** Ken Fujii, Tomo Kondo, Akatsuki Kimura

## Abstract

The centrosome is a major microtubule-organizing center in animal cells. The intracellular positioning of the centrosomes is important for proper cellular function. One of the features of centrosome positioning is the spacing between centrosomes. The spacing activity is mediated by microtubules extending from the centrosomes; however, the underlying mechanisms are not fully understood. To characterize the spacing activity in the *Caenorhabditis elegans* embryo, a genetic setup was developed to produce enucleated embryos. The centrosome duplicated multiple times in the enucleated embryo, which enabled us to characterize the chromosome-independent spacing activity between sister and non-sister centrosome pairs. We knocked down genes in the enucleated embryo and found that the timely spacing was dependent on cytoplasmic dynein. Based on these results, we propose a stoichiometric model of cortical and cytoplasmic pulling forces for the spacing between centrosomes. We also found a dynein-independent but non-muscle myosin II-dependent movement of the centrosomes in a later cell cycle phase. The dynein-dependent spacing mechanisms for positioning the centrosomes revealed in this study is likely functioning in the cell with nucleus and chromosomes, including the processes of centrosome separation and spindle elongation.

## Introduction

Centrosomes are the major microtubule organization centers in animal cells (Azimzadeh and Bornens, 2007). Centrosomes cooperate with microtubules and play important roles in intercellular transport and cell division (O’Connell, 1999; Meraldi, 2016). The positioning of the centrosome in the cell is important for various cellular functions (Tang and Marshall, 2012; Elric and Etienne-Manneville, 2014). In the interphase, centrosomes tend to be located at the cell center. Because the centrosome is associated with the nucleus, this position is important for positioning the nucleus at the cell center (Silkworth et al., 2012). During the mitotic phase, the two centrosomes become the poles of the mitotic spindle and their positions define the direction and asymmetry of cell division (Grill et al., 2001).

The position of centrosomes is controlled by the forces generated by microtubules and motor proteins associated with microtubules. Using microtubules and motor proteins, the centrosome interacts with various structures such as the cell cortex, cytoplasmic vesicles, nucleus, and chromosomes (Dogterom and Yurke, 1997; Grill and Hyman, 2005; Gönczy et al., 1999; Kimura and Kimura, 2011; Malone et al., 2003; Mogilner et al., 2006). In addition to these intercellular structures, the centrosomes interact with each other to position themselves. The two centrosomes sharing a common cytoplasm appear to repel each other. Such repulsive movement is observed along the nuclear surface, which is known as centrosome separation, and in the mitotic spindle. The centrosomes take space between each other, independent of sliding along the nuclear surface or spindle formation. In this study, this nucleus- and spindle-independent activity is defined as the “spacing” activity of the centrosomes. In a classic experiment demonstrating the formation of a cell division furrow between non-sister pairs of centrosomes (i.e., the “Rappaport furrow”), these pairs took space between each other (Oegema and Mitchison, 1997; Rappaport, 1961). In *Drosophila* syncytium cells, the nuclei and spindles are positioned with a certain spacing (Kanesaki et al., 2011; de-Carvalho et al., 2022; Telley et al., 2012). Similar spacing was observed in the oocytes of drug-treated marine ascidians (Khetan et al., 2021), and in the self-organized cell-like organization of *Xenopus* egg extracts (Cheng and Ferrell, 2019). These observations suggest a repulsive interaction between the centrosomes.

The sliding of plus-end-directed motors between antiparallel microtubules that elongate from each pair of centrosomes is an underlying mechanism for the repulsive interaction between the centrosomes (Baker et al., 1993). This model is analogous to the mechanism underlying the spindle pole separation in *Drosophila* anaphase B (Brust-Mascher et al., 2004). In support of this model, bipolar kinesin-5 (Klp61F), a microtubule-bundling protein PRC1 (Fascetto/Foe), and kinesin-4 (Klp3A) were demonstrated to be localized to slide antiparallel microtubules between the centrosomes in the *Drosophila* syncytium (Deshpande et al., 2021). PRC1 (Prc1E) and kinesin-4 (Kif4A) have been shown to separate centrosomes from *Xenopus* egg extracts (Nguyen et al., 2014, 2017). In summary, previous studies on the spacing activity between centrosomes have focused on plus-end-directed motor sliding along antiparallel microtubules. Other mechanisms underlying the spacing between the centrosomes are unknown.

Here, we aimed to reveal a novel mechanism for spacing between centrosomes. The *Caenorhabditis elegans* embryo is a well-studied model system for centrosome biology. Interestingly, unlike humans, *Xenopus,* and *Drosophila*, the *C. elegans* orthologs of proteins involved in the sliding of antiparallel microtubules (BMK-1 [kinesin-5 ortholog], SPD-1 [PRC1 ortholog], and KLP-19 [kinesin-4]), are not required for the elongation of the mitotic spindle in the embryo (Saunders et al., 2007; Lee et al., 2015; Powers et al., 2004). The investigation of the spacing activity of the centrosomes in the *C. elegans* embryo should have an impact on *C. elegans* biology and also on other species, because kinesin-independent spacing has been suggested in other species (Donoughe et al., 2022).

It is challenging to characterize centrosome spacing activity, which is independent of the nucleus and spindle, in the *C. elegans* embryo. The centrosomes in the wild-type *C. elegans* embryo are always associated with the nucleus or spindle, and embryonic cells do not form syncytia. Inactivation of the *zyg-12* gene offers some information on centrosome spacing, independent of the nucleus, as this gene encodes a KASH domain protein essential for the association between the centrosome and the nucleus (Malone et al., 2003). In *zyg-12*-impaired cells, centrosomes move even when they are not attached to the nucleus until they are incorporated into the mitotic spindle. Upon inactivation of *zyg-12*, the two centrosomes in the 1-cell stage embryo separate, indicating that spacing activity independent of the nucleus exists. Centrosome separation is also impaired by the RNAi of genes involved in the cortical pulling force, suggesting that this force affects spacing (De Simone et al., 2016). However, it is unclear how the two centrosomes move in opposite directions, instead of being pulled toward the same cortical region. It has been proposed that cytoplasmic flow contributes to this process (De Simone et al., 2016). However, this model does not ensure that the two centrosomes move toward the opposite directions. In addition, this cytoplasmic flow occurs only in the 1-cell stage. Therefore, this mechanism cannot be considered a general mechanism for centrosome spacing. *zyg-12* affects the interaction between the nucleus and centrosome, but not spindle formation. Therefore, a spacing mechanism independent of the spindle could not be identified in *zyg-12*-impaired cells. Spacing activity, which is independent of nuclei and spindles and is general to multiple stages of embryogenesis, was expected but uncharacterized in the *C. elegans* embryo.

## Results

### Establishment of enucleated *C. elegans* embryos by genetic manipulation

To characterize the spacing activity between the two centrosomes, an experiment was designed to remove the chromosomes from the *C. elegans* embryo (“enucleated embryo”). Enucleated *C. elegans* embryos were produced in classic experiments by Schierenberg and Wood, where the nuclei were removed by penetration of the eggshell by laser microsurgery, followed by pressing the cytoplasm to push the nucleus out of the eggshell (Schierenberg and Wood, 1985). This method often removes the centrosome together with the nuclei and is unsuitable for analyzing the behavior of the centrosome.

To create an enucleated embryo with centrosomes, the paternal and maternal chromosomes were removed by using *emb-27* mutant sperm (Sadler and Shakes, 2000), and by knocking down the *klp-18* gene (Segbert et al., 2003) (Fig. S1). *emb-27* encodes a subunit of the anaphase-promoting complex. Its mutation causes chromosome segregation defects and produces centriole-containing fertilization-competent enucleated sperm (Sadler and Shakes, 2000; Kondo and Kimura, 2018). *klp-18* is a member of the kinesin family and is required for oocyte meiosis. The *klp-18* knockdown oocyte occasionally extrudes all chromosomes into the polar body, resulting in embryos without maternal chromosomes. By mating the worms with *emb-27* mutant sperms and *klp-18* knockdown oocytes, we expected to obtain enucleated embryos to characterize the spacing activity of the centrosomes, independent of the nucleus and spindle.

In this study, *C. elegans* strains were used in which the centrosomes (γ-tubulin), chromosomes (histone H2B), and cell membranes (PH^PLCδ1^) were visualized using green fluorescent protein (GFP) (Fig. 1 and Table S1). In control 1-cell stage embryos, sperm- and oocyte-derived pronuclei appeared after fertilization (Fig. 1A and Movie S1). The two centrosomes associated with the sperm pronucleus moved toward the cell center and met the oocyte pronucleus before the first cytokinesis. When *emb-27* (*g48*ts) males were mated with control hermaphrodites, the sperm pronucleus was absent, as reported previously (Sadler and Shakes, 2000) (Fig. 1B and Movie S2). The centrosomes migrated toward the cell center and met the oocyte pronucleus. The *emb-27* mutants affected the number of centrosomes supplied by the sperm (Kondo and Kimura, 2018, 2019). Consequently, the presence of 1-4 centrosomes was observed in the 1-cell stage *emb-27* mutant and in the enucleated embryo. Oocyte pronuclei were not detected in the *klp-18* (RNAi) embryos, as reported previously (Segbert et al., 2003) (Fig. 1C and Movie S3). We designed an experiment to obtain embryos without chromosomes by mating *emb-27* (*g48*ts) males with *klp-18* (RNAi) hermaphrodites. Enucleated embryos were successfully obtained using this experimental setup (Fig. 1D and Movie S4). No sperm-, oocyte-derived pronuclei, or chromosomes were detected inside the embryonic cells at a subsequent stage. The chromosome signals of the polar bodies were detected outside the cytoplasm.

**Fig. 1:**
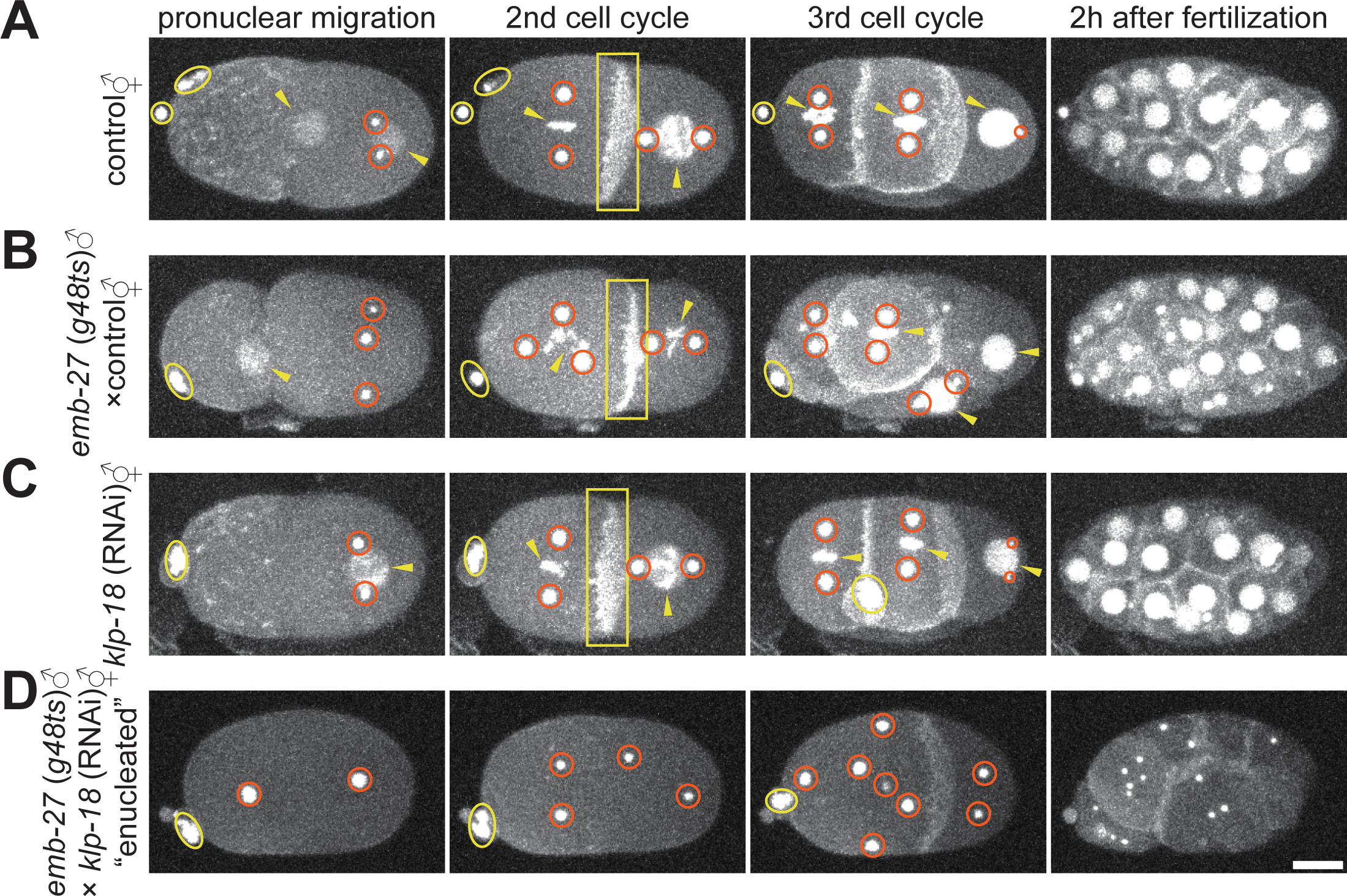
Establishment of enucleated *C. elegans* embryos by genetic manipulation. (A-D) γ-tubulin (centrosome), histone-H2B (chromosome), and PH ^PLCδ1^ (cell membrane) were labeled with GFP. Red circles indicate centrosomes. Yellow arrowheads indicate the pronucleus, nucleus, or chromosome. The yellow oval indicates the polar body. Yellow squares indicate the cell membranes. z-maximum projections. Scale bar, 10 μm. (A) A time lapse imaging series of an embryo of the control strain (CAL0181) grown at 16 °C, with imaging at 18–22 °C. The reproducibility of the observations was confirmed (n = 5). (B) An embryo from a hermaphrodite of CAL0181 strain mated with males of CAL0051 strain. Both strains were grown at 25 °C, with imaging at 18–22 °C (n = 5). The embryo of this figure initially possessed three centrosomes at the 1-cell stage. (C) An embryo of the hermaphrodite of CAL0181 with *klp-18* (RNAi) grown at 25 °C after injection, with imaging at 18–22 °C. (n = 5). (D) An embryo of the hermaphrodite CAL0181 with *klp-18* (RNAi) was mated with males of the CAL0051 strain. Both strains were grown at 25 °C, with imaging at 18– 22 °C. (n = 7).

In the enucleated embryo, the centrosomes moved dynamically, which is the main topic of this study. This indicated that centrosomes can move without requiring nuclei or chromosomes. Interestingly, the centrosomes were duplicated periodically for multiple rounds, which appeared to correspond to the cell cycle. Cytokinesis was impaired, at least for the several cell cycles (Fig. 1D), possibly because of chromosomal loss (Bringmann and Hyman, 2005). Most importantly for this study, the positions of these centrosomes did not overlap but were spread throughout the cell (Fig. 1D), suggesting the existence of the spacing activity. Therefore, the enucleated *C. elegans* embryo is suitable for analyzing the centrosome spacing mechanism, which is independent of the nucleus and spindle. In summary, sister and non-sister centrosomes shared a common cytoplasm and moved dynamically in the enucleated embryo.

### A repulsive spacing between sister and non-sister centrosomes was observed in the enucleated embryo

To characterize the force acting between centrosomes, the change in the distance between centrosomes over time was quantified. In the present study, we focused on the time window corresponding to the 2-cell stage in control embryos with nuclei (Fig. 2A and 2B). In the enucleated embryo, the cytokinesis failed; therefore, the cytoplasm did not divide into two in the “2-cell stage.” At this stage, four (or more, depending on the number of centrosomes in the 1-cell stage, as explained in the previous section) centrosomes of two sister and non-sister pairs coexisted in the common cytoplasm. We focused on this stage because it was the earliest stage at which potential interactions between non-sister pairs of centrosomes can be tracked. We set the time zero of the time window when we first detected two discrete centrosome (γ-tubulin::GFP) spots for the sister pair after the 2nd centrosome duplication (Fig. 2A). The time window ended when the signal of the spot became too weak to be identified, or when the spot was duplicated into two in the subsequent round of centrosome duplication.

**Fig. 2:**
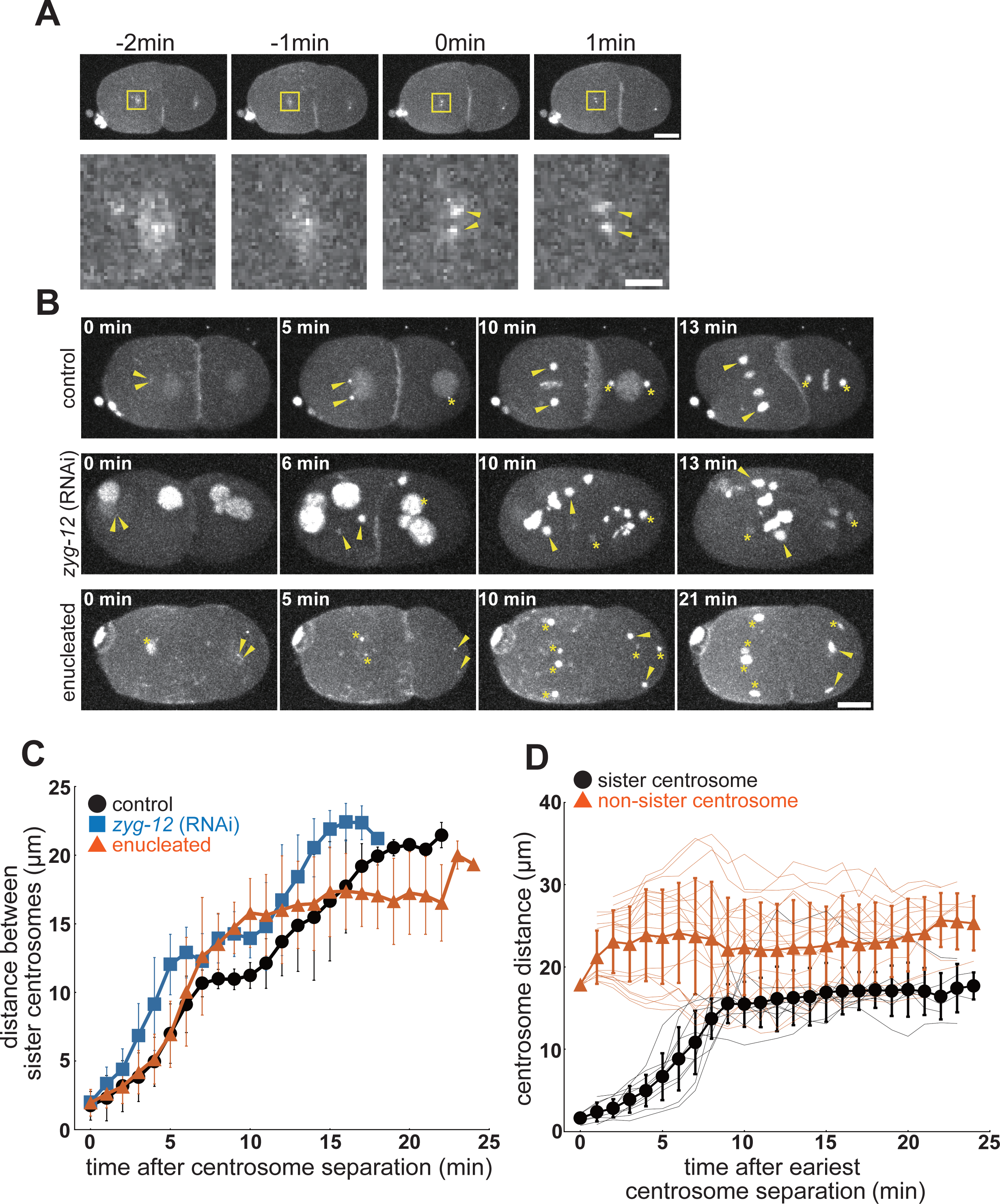
Characterization of centrosome dynamics during the 2-cell stage. (A) The definition of time zero of the 2-cell stage. Representative time series images (upper) and the enlarged images of the yellow box (lower) of an enucleated embryo. The time point when we detected two discrete spots in the cloud of the γ-tubulin::GFP signal was defined as time zero. The yellow arrowheads indicate two discrete spots of centrosomes. z-maximum projections. Scale bars represent 10 μm for upper panels and 2 μm for lower panels. (B) A time lapse imaging series of an embryo of the control strain (CAL0181), a DE90 strain with *zyg-12* (RNAi) embryo, and an enucleated embryo in the 2-cell stage. The yellow arrowheads indicate a pair of sister centrosomes. The asterisks indicate the other centrosomes in the images. z-maximum projections. Scale bar, 10 μm. (C) The quantification of the distance between sister centrosomes. The mean and standard deviation (S.D.) are shown with the symbol and the error bar, respectively. Black circle, control embryos (8 sister pairs from 5 embryos). Blue square, *zyg-12* (RNAi) embryos (7 sister pairs from 5 embryos). Red triangle, enucleated embryos (10 sister pairs from 5 embryos). (D) Distance between the sister- and non-sister-pairs of centrosomes in the enucleated embryo. The distances between the non-sister pairs are calculated for all possible pairs of the non-sisters. Individual samples are shown with thin lines. To compare the sister- and non-sister pairs, the time after the earliest centrosome separation of the cell is indicated in the horizontal axis, which is slightly different from the time in (C). The mean and S.D. are shown with the symbol and the error bar, respectively. Black circle, sister pairs (10 pairs from 5 embryos). Red triangle, non-sister pairs (20 pairs from 5 embryos).

The distance between sister pairs of centrosomes in the enucleated embryos, controls (i.e., embryos with nuclei), and *zyg-12* (RNAi) embryos was compared (Fig. 2B and 2C). At early time points in the time window in the control embryos, sister centrosomes slid along the nuclear surface to position themselves at opposite poles of the nucleus (Fig. 2C black, and Movie S5), as reported previously (Gönczy et al., 1999). In the enucleated embryo, the sister centrosomes separated at a speed similar to that in the control embryos, indicating that the spacing was independent of the nucleus (Fig. 2C red, and Movie S7). The nucleus-independent spacing was consistent with previous observations for *zyg-12* (RNAi), in which centrosomes were not associated with nuclei (Malone et al., 2003) (Fig. 2C blue, and Movie S6). At later time points, for enucleated embryos, unlike the control embryos, the separation of sister centrosomes did not pause at the distance of the nuclear diameter (∼10 µm), but continued to increase. This behavior can be explained by the loss of association with the nucleus. In *zyg-12* (RNAi) embryos, the separation of the centrosomes slowed as the centrosomes formed the mitotic spindle until the centrosomes separated again in anaphase. In conclusion, centrosomes have the intrinsic ability to separate from their sister centrosomes, independent of their sliding activity along the nuclear surface. In control embryos, the nucleus tethered the sister centrosomes. Therefore, the centrosomes did not separate further until nuclear envelope breakdown (NEBD).

An advantage of enucleated embryos is that the interaction between non-sister centrosomes that share the cytoplasm can be characterized. Notably, the distance between non-sister centrosomes was always longer than that between sister centrosomes (Fig. 2D and S2). Even though the centrosomes moved dynamically within the embryo, the distances between the non-sisters did not become shorter than the minimal distance between sister pairs at each time point. The results indicated that similar spacing activity existed between sister and non-sister pairs of centrosomes. Therefore, repulsive spacing activity is intrinsic to the centrosome.

### Dynein-dependent pulling forces were responsible for the timely spacing activity

To obtain insights into the mechanism of centrosome spacing, we searched for genes involved in this activity in enucleated embryos. Cortical pulling forces, which pull microtubules from force generators located in the cell cortex, contribute to centrosome separation in *Drosophila* (Cytrynbaum et al., 2003). In *C. elegans*, knockdown of genes required to generate the cortical pulling force (e.g., *gpr-1/2* (RNAi)) impaired spacing in *zyg-12* knockdown embryos (De Simone et al., 2016). We knocked down *gpr-1/2* in an enucleated embryo to inhibit the cortical pulling force, and found the distance between the centrosomes was shortened (Fig. 3A, 3B orange, and Movie S8). A significant difference (*p*<0.01 at 10-min, Wilcoxon rank sum test) was found between enucleated embryos and enucleated embryos with *gpr-1/2* (RNAi) in the distance between sister centrosomes. The distances for enucleated embryos and enucleated embryos with *gpr-1/2* (RNAi) were 15.8 ± 2.5 μm and 6.9 ± 3.2 μm (mean ± SD), n=10 and 13, respectively, at the 10-min timing when the distance between centrosomes in enucleated embryos reached near saturation. The results indicated that the cortical pulling force mediated spacing activity. Centrosomes were separated to some extent in enucleated *gpr-1/2* (RNAi) embryos. This result was consistent with the incomplete separation of *zyg-12*; *gpr-1/2* (RNAi) embryos (De Simone et al., 2016). While we could not exclude the possibility that the cortical pulling force was not completely impaired by *gpr-1/2* (RNAi), we expected that different factors were involved in centrosome spacing activity based on the following observations.

**Fig. 3:**
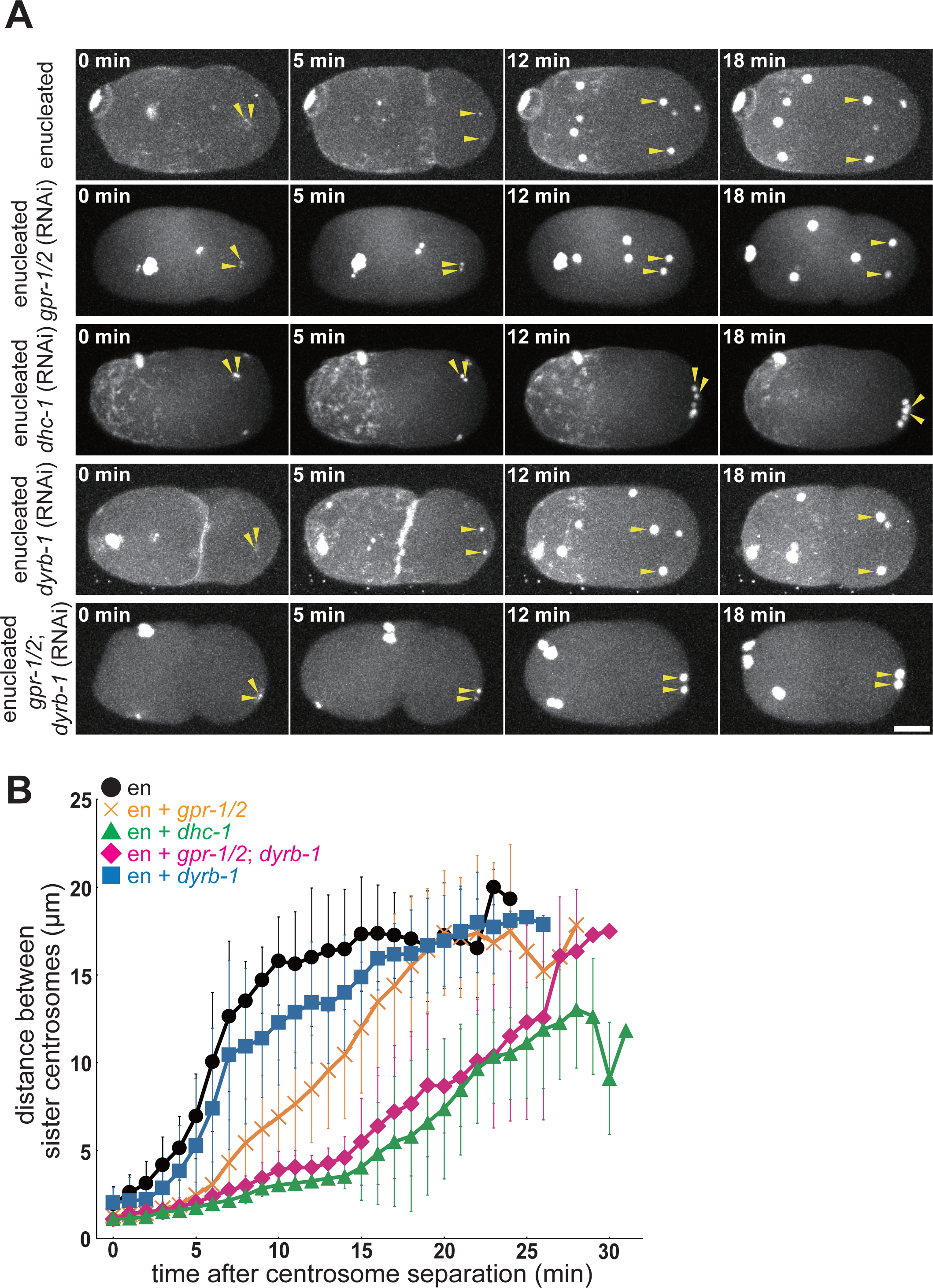
Centrosome spacing activity depends on cortical- and cytoplasmic-pulling forces. (A) Time lapse imaging series of an embryo of an enucleated embryo, and *gpr-1/2* (RNAi), *dhc-1* (RNAi), *gpr-1/2*; *dyrb-1* (RNAi), and *dyrb-1* (RNAi) enucleated embryos. The yellow arrowheads indicate a pair of sister centrosomes. z-maximum projections. The time zero is when two discrete centrosome (γ-tubulin) spots were detected for a sister pair of interest after the 2nd centrosome duplication (Described in Fig. 2A). Scale bar, 10 μm. (B) The quantification of the distance between sister centrosomes. The mean and S.D. are shown with the symbol and the error bar, respectively. Black circle, enucleated embryos (10 sister pairs from 5 embryos). Orange cross, *gpr-1/2* (RNAi) enucleated embryos (13 sister pairs from 5 embryos). Green triangle, *dhc-1* (RNAi) enucleated embryos (12 sister pairs from 5 embryos). Magenta diamond, *gpr-1/2*; *dyrb-1* (RNAi) enucleated embryos (12 sister pairs from 5 embryos). Blue square, *dyrb-1* (RNAi) enucleated embryos (11 sister pairs from 5 embryos).

When *dhc-1* was knocked down in the enucleated embryo, the spacing was almost completely blocked for approximately 20 min, which corresponded to the duration of the cell cycle in the control embryo (Fig. 3A, 3B green, and Movie S9). *dhc-1* encodes the heavy chain subunit of cytoplasmic dynein and is responsible for all microtubule-pulling forces in *C. elegans* embryos (Gönczy et al., 1999; Torisawa and Kimura, 2020). A significant difference was found between enucleated embryos with *gpr-1/2* (RNAi) and enucleated embryos with *dhc-1* (RNAi) (*p*<0.01 at 10-min, Wilcoxon rank sum test). The distance for enucleated embryos with *dhc-1* (RNAi) was 3.1 ± 0.4 μm, n=12. Here, we focused on the near-complete block of spacing for the first ∼20 min under *dhc-1* (RNAi) conditions. Notably, we observed apparent movement of the centrosomes after 20 min. The mechanism of the latter movement is investigated in the final section of this paper. Dynein inhibition impaired the timely spacing activity, which should take place almost completely within 20 min. Therefore, these results suggested that factors other than the cortical pulling force, but are dependent on dynein, contribute to the spacing.

The cytoplasmic pulling force depends on *dhc-1* but not on *gpr-1/2* and drives the centration of the centrosomes and pronuclei (Kimura and Onami, 2005, 2007; Kimura and Kimura, 2011). We expected that the cytoplasmic pulling force would contribute to this spacing. To test this possibility, the *dyrb-1* and *gpr-1/2* genes were simultaneously knocked down in the enucleated embryo. *dyrb-1* encodes a roadblock subunit of the dynein complex, which is not essential for the motor activity of dynein but is required for organelle transport and centration of the centrosome; therefore, it is necessary for the cytoplasmic pulling force (Kimura and Kimura, 2011). In *gpr-1/2*; *dyrb-1* (RNAi)-enucleated embryos, in which the cytoplasmic- and cortical-pulling forces were impaired, centrosome spacing was severely defective, as observed in *dhc-1* (RNAi)-enucleated embryos (Fig. 3A, 3B magenta, and Movie S10). A significant difference was found between the enucleated embryos with *gpr-1/2* (RNAi) and enucleated embryos with *gpr-1/2*; *dyrb-1* (RNAi) in the distance between the sister centrosomes (*p*<0.01 at 10-min, Wilcoxon rank sum test). The distance for enucleated embryos with *gpr-1/2*; *dyrb-1* (RNAi) was 3.9 ± 1.1 μm, n=12. The enucleated embryos with *dyrb-1* (RNAi) repressed the spacing compared to the control (*p*<0.05 at 10-min, Wilcoxon rank sum test), but not as severe as *gpr-1/2; dyrb-1* (RNAi) (Fig. 3A, 3B blue, and Movie S11). The distance for enucleated embryos with *dyrb-1* (RNAi) was 12.3 ± 3.5 μm, n=11. These results showed that cortical and cytoplasmic pulling forces were sufficient to provide spacing between the centrosomes that occurred in the initial 20 min of the 2-cell stage.

### In search for a dynein-dependent mechanism for the spacing between centrosomes

In humans, *Drosophila*, and *Xenopus*, plus-end-directed motors are involved in centrosome spacing for the mitotic spindle (see Introduction) and are considered to be involved in chromosome-independent spacing by acting on antiparallel microtubules emanating from the two centrosomes (Deshpande et al., 2021; de-Carvalho et al., 2022). In contrast, in this study, the minus-end-directed motor dynein provided the necessary force for centrosome spacing in *C. elegans* embryos. Therefore, we aimed to determine how pulling forces mediate the repulsive interactions between centrosomes.

Analogous to the finding of the antiparallel pushing mechanism for spacing activity in *Drosophila* and *Xenopus* from the mechanisms for spindle elongation, we speculated whether we could obtain a clue for the spacing mechanism in *C. elegans* from the mechanisms proposed for spindle elongation in the species. Spindle elongation in *C. elegans* is dependent on dynein. Most of the proposed models for spindle elongation in the *C. elegans* embryo assume that the distribution of microtubules is different between the two centrosomes (Grill et al., 2001; Hara and Kimura, 2009). Farhadifar et al. proposed a mechanism called “the stoichiometric model of cortical pulling forces,” for the spindle elongation in the *C. elegans* embryo that is independent on the distribution of microtubules (Farhadifar et al., 2020). In this model, the two centrosomes of the spindle poles compete for force generators in the cell cortex to be pulled. “Stoichiometric” means that one force generator can pull only one centrosome, which is located the nearest. This model ensures that anterior and posterior cortexes pull only the anterior and posterior centrosomes, respectively. Here, we applied a stoichiometric model to explain the spacing activity of four or more centrosomes in the enucleated embryo.

### Quantification of the length distribution of the microtubules in the *C. elegans* embryo

The original stoichiometric model (Farhadifar et al., 2020) assumed long and stable microtubules (i.e., exponential decay with a characteristic length of 20 μm). The length distribution of the microtubules should be critical for the stoichiometric models, and thus we quantified the length distribution of the microtubules experimentally (Fig. 4 and S3). We assumed that the brightness intensity of β-tubulin::GFP signal above its cytoplasmic average was proportional to the number of microtubules and quantified the value (Fig. 4A and 4B). The quantified signal intensity fitted well with a Weibull distribution of *S*(*l*) = *S*_0_×EXP[-{(*l-l*_0_)/*ξ*}^*P*], where *S*(*l*) is the signal intensity of microtubules with their length over *l, l*_0_ is the size of the centrosome, *S*_0_ is the intensity at the surface of the centrosome, *ξ* is the length-scale, and *P* is a parameter for how the distribution is affected by the length (Fig. 4B and see Materials and Methods). The estimated distribution of microtubule lengths did not change dramatically during the observation period (Fig. 4C and S3) or among different samples (Fig. S3). Therefore, we calculated the average distribution of all samples at all the time points (Fig. 4C, 4D and S3). Our fitting of the average distribution to the Weibull distribution revealed *l*_0_ = 1.6 μm, *ξ* = 2.3 μm, and *P* = 0.79.

**Fig. 4:**
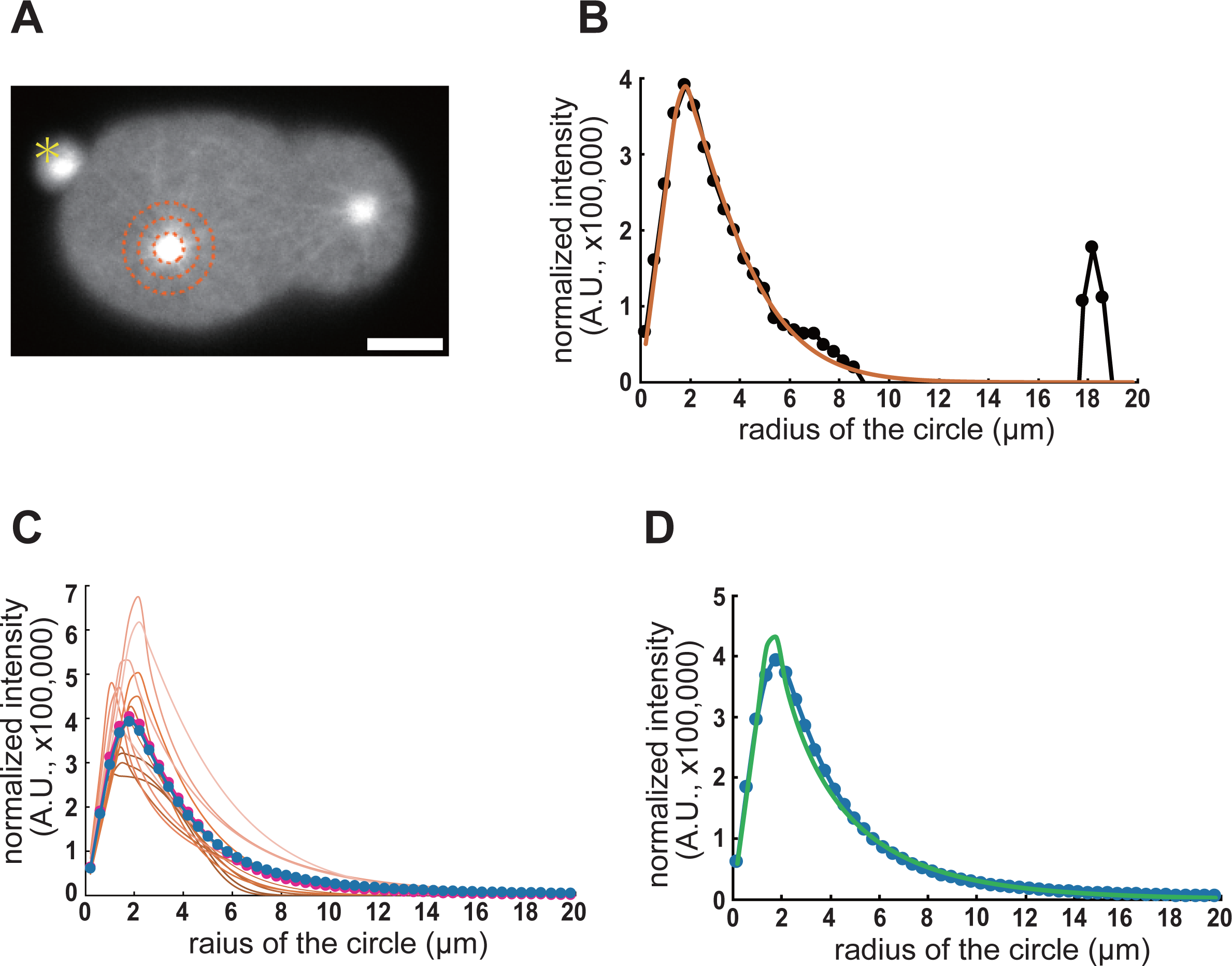
Distribution of the microtubule length in enucleated embryos. (A) A representative image of the β-tubulin signal in enucleated *C. elegans* embryos. The brown dot circles indicate areas 2 μm, 4 μm, and 6 μm from center of the aster. The yellow asterisk indicates the polar body. Scale bar, 10 μm. (B) The fitting analysis result of the distribution of the β-tubulin signal in (A). The subtracted intensity value (see Materials and Methods for the details) is shown with the black circle and line. Brown line is the fitted curve. (C) The fitting results of the β-tubulin signals in an enucleated embryo. Fitted curves for each time point are shown in brown lines. Darker colors indicate earlier time points. Lighter colors indicate later time points. The average fitting curve is shown with magenta dots and lines. The average fitting curve from 5 embryos is shown with blue dots and lines. (D) The average fitting curve from 5 embryos is shown with blue dots and lines. The fitting of the average fitting curve is shown with a green line. The value from this result is applied for simulation (Table S2).

### The stoichiometric model of cortical and cytoplasmic pulling forces as a mechanism for the repulsive spacing between centrosomes in the *C. elegans* embryo

Our present analyses revealed that both cortical and cytoplasmic pulling forces act for the spacing between the centrosomes (Fig. 3). Therefore, we added the cytoplasmic pulling forces (Fig. 5A, 5B and Table S2) to the stoichiometric model of the cortical pulling force by Farhadifar et al. (2020). This modified version of the stoichiometric model of cortical and cytoplasmic pulling forces reproduced the major features of our experimental measurements (Fig. 5C and 5D). In the 3-dimensional space (ellipsoid), we placed force generators in the cytoplasm and in a thin layer of the cortex uniformly (Fig. 5B), similar to our previous simulation (Kondo and Kimura, 2019). The centrosomes were positioned corresponding to their initial positions in representative experiments (Table S3). The simulation was conducted by iterating the processes of microtubule growth, summing the pulling forces calculated as in the original stoichiometric model (Farhadifar et al., 2020) but adding the contribution from the cytoplasmic force generators, and moving the centrosomes.

**Fig. 5:**
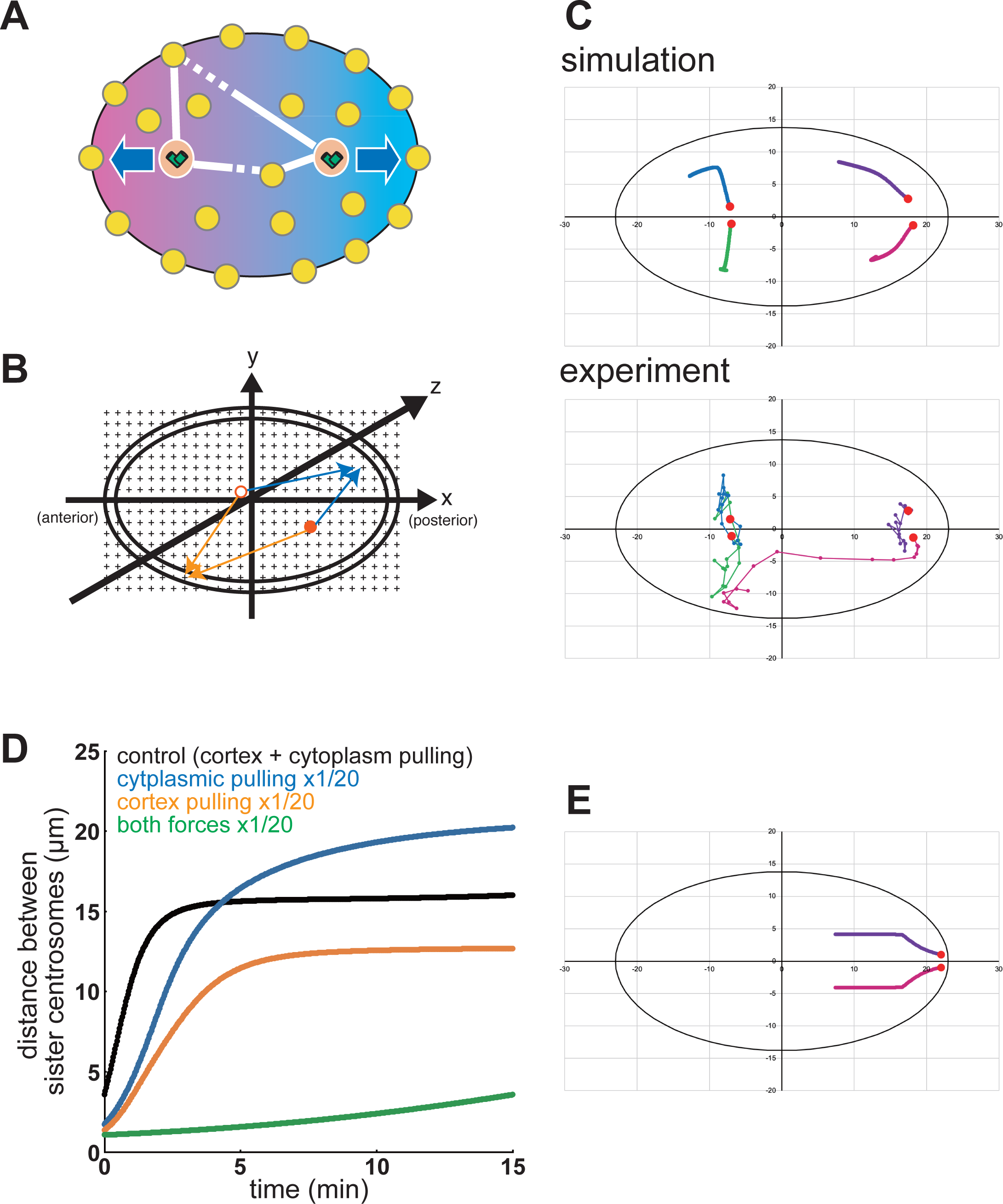
The stoichiometric model of cortical and cytoplasmic pulling forces. (A) Schematic of the model. Each centrosome (orange circle with two cylinders) is pulled by force generators (yellow circles) at the cell cortex and the cytoplasm. One force generator can pull only one centrosome, which is the nearest. The centrosome connected by solid lines are pulled by the force generator. (B) Schematic of the simulation setup. The ellipsoids represent the cell (the outer layer: the cortex, the inner mass: the cytoplasm). Red circles (open and filled) are the centrosomes. Black crosses on the lattice are the force generators evenly distributed. The force generators at the cortex, or the cytoplasm, pull the centrosomes depending on the distance between the force generator and each centrosome (orange or blue arrows, respectively). (C) Trajectories of the centrosomes in a representative enucleated embryo (lower) and the simulation with the same initial positions of the 4 centrosomes (upper). The initial positions of the centrosomes are shown with red circles. The trajectories of the same color indicate the same initial positions. (D) Simulated distance between the sister centrosomes in the simulation shown in C (black line), and in simulations with reduced (5%) cortical pulling forces (orange line), with reduced (5%) cytoplasmic pulling forces (blue line), and with a condition where the both pulling forces are reduced (5% for each) (green line). (E) Simulation for the separation and migration of the centrosomes in the pronuclear migration stage in the wild-type (intact nucleus). The trajectories of the two centrosomes are shown in magenta and purple lines. The initial positions of the two centrosomes are set near the posterior end of the embryo, and the initial spacing between the centrosomes is 2 μm. The intact nucleus was simulated by restricting the distance between the two centrosomes not exceeding the nuclear diameter (8 μm).

For the simulation parameter values, we followed the values of the original stoichiometric model for cortical pulling forces (Farhadifar et al., 2020). See the Materials and Methods and Table S2 for details on the parameter values. The simulations (Fig. 5D) reproduced centrosome spacing of similar magnitudes and increased rates for control, *dyrb-1* (RNAi), *gpr-1/2* (RNAi), and *dhc-1* (RNAi) enucleated embryos shown in Fig. 3B. The trajectories of the centrosomes inside the cell were also similar in the simulations and experiments (Fig. 5C and S4). The trajectories fluctuated more in the experiments, possibly because of random fluctuations in the cytoplasm (Guo et al., 2014). Overall, the numerical simulation results supported the feasibility of the stoichiometric model of the cortical and cytoplasmic pulling forces.

In addition, the stoichiometric model of cortical and cytoplasmic pulling forces reproduced the separation and centration of centrosomes (Fig. 5E) associated with the sperm-derived pronucleus of the 1-cell stage embryo (Albertson, 1984; Gönczy et al., 1999). For the simulation with the nucleus, the two centrosomes were connected with an elastic bar with the length of the nuclear diameter (8 μm), by which the centrosomes attract each other when the distance between them exceeds the nuclear diameter. This result supports the feasibility of the model even for cells with nuclei. In addition, the application of the distribution to the original stoichiometric model resulted in the elongation of the spindle for almost as long as the cell length (Fig. S5). This is likely because the microtubules are shorter than the assumed distribution and the original model assumes only the cortical pulling force. The addition of cytoplasmic pulling forces to the original stoichiometric model enabled elongation to a reasonable extent (Fig. S5), suggesting that the stoichiometric model of cortical and cytoplasmic pulling forces accounts for spindle elongation, in addition to the separation and centration of centrosomes, in normal embryos with nuclei and chromosomes.

### Myosin-dependent movements of the centrosomes in the *C. elegans* embryo

In this study, a large movement of the centrosomes was found approximately 20 min after detecting two centrosomes in *dhc-1* (RNAi)-enucleated embryos (Fig. 3B green, Movie S9). In *dhc-1* (RNAi) embryos with nuclei, the centrosomes did not separate during interphase in the 1-cell stage, and a small spindle-like structure was formed near the cortex, indicating that dynein was responsible for all centrosome movements until the spindle formation stage in normal embryos (Fig. 6A). We noticed that the centrosomes moved over a large distance in *dhc-1* (RNAi) embryos with nuclei in a later cell cycle phase, indicating that large centrosome movement was not specific to *dhc-1* (RNAi)-enucleated embryos (Movie S9 and S12).

**Fig. 6:**
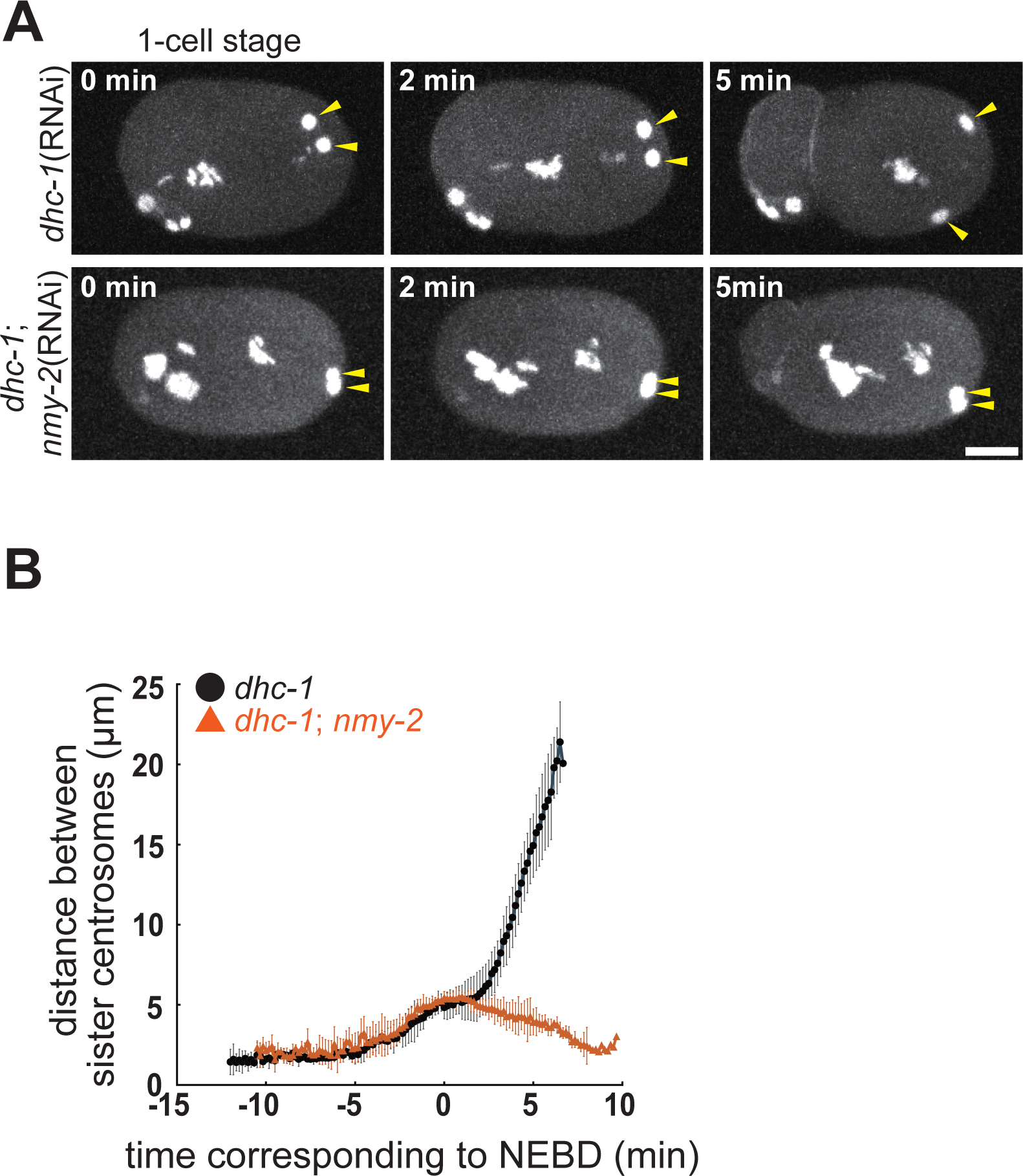
The myosin-dependent movement of the centrosomes. (A) Time lapse imaging series of a *dhc-1* (RNAi) embryo, and a *dhc-1*; *nmy-2* (RNAi) embryo of the DE90 strain at 1-cell stage. GFP labeled γ-tubulin (centrosome), histone-H2B (chromosome), and PH ^hPLCIIIδ1^ (cell membrane) are shown. The yellow arrowheads indicate a pair of sister centrosomes. z-maximum projections. Scale bar, 10 μm. (B) The quantification of the distance between sister centrosomes. The mean and S.D. are shown by the symbol and error bar, respectively. Black circle, *dhc-1* (RNAi) embryos (5 sister pairs from 5 embryos). Red triangle, *dhc-1*; *nmy-2* (RNAi) embryos (5 sister pairs from 5 embryos).

A large movement occurred near the time of cytokinesis (in *dhc-1* (RNAi) embryos); therefore, we speculated the involvement of actomyosin, which drives the constriction of the contractile ring and the accompanying cytoplasmic flow toward the equatorial plane (Bray and White, 1988; Khaliullin et al., 2018). Although cytokinesis does not occur at this stage in enucleated embryos, cytoplasmic flow may move the centrosomes. Unfortunately, an enucleated embryo could not be obtained upon the knockdown of *nmy-2*, which encodes non-muscle myosin II and is required for cytoplasmic flow (Munro et al., 2004). This was possibly because NMY-2 was required for polar body extrusion (Dorn et al., 2010) and lowered oocyte enucleation efficiency. To this end, *dhc-1* and *nmy-2* were simultaneously knocked down in nucleated embryos. We observed an impairment in dynein-independent centrosome movements, indicating that a large movement is driven by cytoplasmic flow (Fig. 6A, 6B and Movie S13). Differences in the distance between centrosomes 5 min after NEBD were tested using the Wilcoxon rank sum test between *dhc-1* (RNAi) and *dhc-1*; *nmy-2* (RNAi) experiments (*p*<0.01). The mean values for *dhc-1* (RNAi) and *dhc-1*; *nmy-2* (RNAi) were 14.9 ± 2.0 μm and 3.9 ± 0.9 μm, n=6 and 5, respectively. In conclusion, cytoplasmic flow drives the large movement of centrosomes during late mitosis, which occurs when dynein is inhibited.

## Discussion

### Enucleation of *C. elegans* embryos

Chromosomes are essential for cellular function because they carry genetic information and constitute a core component of intracellular organelles. Before NEBD, the chromosome forms the cell nucleus, whereas after NEBD, it creates a mitotic spindle. Historically, chromosome removal from cells has enabled important modeling in cell biology studies (Goldman and Pollack, 1974). In addition to the mechanical removal of the nucleus (e.g. by centrifugation or microneedles), genetic manipulation can also be used to investigate the function of nuclei. The *gnu* mutant of *Drosophila* revealed a semi-enucleated system in *Drosophila* embryos in which the nucleus does not divide but forms one giant nucleus, while the centrosomes continually duplicate and separate (Freeman et al., 1986). In the *gnu* mutant, the separation of the centrosomes is almost entirely independent of the existence of the nuclei (de-Carvalho et al., 2022). In the present study, we established a genetic method to obtain enucleated *C. elegans* embryos by combining previously established methods to remove chromosomes from sperm (Sadler and Shakes, 2000) and oocytes (Segbert et al., 2003) (Fig. S1). Our established method produces enucleated embryos in a non-invasive manner, whereas classical enucleated *C. elegans* embryos must be obtained by penetrating the eggshell using laser microsurgery (Schierenberg and Wood, 1985). Unlike the *Drosophila gnu* mutant, our method completely removed chromosomes from the embryo. We expect our established method to be applied in various studies and not limited to centrosome research because *C. elegans* embryos are widely used model organisms.

### Chromosome-independent and dynein-dependent spacing between the centrosomes

Using the enucleated *C. elegans* embryo, we demonstrated that spacing activity, independent of the chromosome, existed in the *C. elegans* embryo before and after NEBD and between the sister and non-sister-centrosomes. Nucleus-independent spacing between sister centrosomes before NEBD has been previously observed in the *zyg-12* mutant, in which the association between the nucleus and centrosomes is impaired (Malone et al., 2003; De Simone et al., 2016). In contrast, as the mitotic spindle forms and most cells divide in *zyg-12* mutants, the chromosome-independent interaction after NEBD and that for non-sister centrosomes could not be previously addressed. In addition, even in the *zyg-12* mutants, nuclei remain in the cytoplasm between the centrosomes during interphase, which acts as an obstacle to microtubule growth in these regions. Therefore, the enucleated embryo is a good model system for studying the interaction between centrosomes in an intracellular space without physical obstacles.

We demonstrated that spacing between sister and non-sister centrosomes until cytokinesis was completely impaired the knockdown of dynein (*dhc-1*). This result indicated that kinesin-dependent pushing between the centrosomes, as revealed in *Drosophila* and *Xenopus* (Telley et al., 2012; Nguyen et al., 2017, 2014), did not occur in *C. elegans* embryo. Our observation in *C. elegans* is consistent with previous research showing that the centrosomes rarely moved in *dhc-1* (RNAi) embryos (Gönczy et al., 1999) and that the molecules involved in pushing did not impair the elongation of the mitotic spindle in the *C. elegans* embryo (Saunders et al., 2007; Powers et al., 2004; Lee et al., 2015). Our study using enucleated embryos demonstrates the direct requirement of dynein (*dhc-1*) for the centrosome spacing.

### The stoichiometric model of cortical and cytoplasmic pulling forces

The major involvement of dynein indicates that centrosome spacing in *C. elegans* is driven by a pulling force outside the centrosome pairs. The separation of centrosomes by outward pulling forces commonly occurs before NEBD (Cytrynbaum et al., 2003; Gönczy et al., 1999; De Simone et al., 2016) or spindle elongation (anaphase B) (Grill et al., 2001). However, it has not yet been determined why the two adjacent centrosomes are pulled toward opposite directions. In mitotic spindles, the spindle itself exhibits bipolarity, and this difference is established upon spindle formation. In other cases, the cell nucleus may amplify the asymmetry by positioning itself between the centrosomes and obstructing the growth of microtubules toward the nucleus (Donoughe et al., 2022). Cortical flow has also been proposed to separate centrosomes (De Simone et al., 2016); however, the mechanism that ensures centrosome movement toward the opposite direction has not been clarified.

Here, we extend the idea of the stoichiometric model of cortical pulling forces proposed by Farhadifar et al. (2020) for spindle elongation to explain the spacing activity independent of the nuclei and spindle. An important modification is the addition of a cytoplasmic pulling force. This was required to explain the RNAi phenotypes (Fig. 3) and spindle elongation with the experimentally obtained distribution of microtubule length (Fig. 4 and S5). A stoichiometric model was proposed as the underlying mechanism of spindle elongation (Farhadifar et al., 2020). The idea that a force generator can pull only one microtubule among multiple microtubules, potentially reaching the force generator, is in line with the previously proposed force-generator-limited model (Grill et al., 2003; Grill and Hyman, 2005). We noticed that during spindle elongation, the microtubules from the two spindle poles rarely overlap (Tada, KF, AK, Funahashi et al., submitted). This observation indicated that the competition assumed in the stoichiometric model may not be critical for spindle elongation. In contrast, in the case of enucleated embryos, there was no apparent bias in the direction of microtubule elongation. We demonstrated that a stoichiometric model of cortical and cytoplasmic pulling forces is critical for centrosome spacing in *C. elegans* embryos. Because we demonstrated the existence of spacing activity and a stoichiometric model of cortical and cytoplasmic pulling forces as the underlying mechanism, the model ensures spindle elongation even without the formation of mitotic spindles (Fig. S5).

We showed that the stoichiometric model of cortical and cytoplasmic pulling forces corroborated the spacing dynamics of centrosome pairs in control and gene-knockdown enucleated embryos (Fig. 5 and S4). Moreover, the model explained the separation and centering of the normal embryo with the nucleus when centrosomes were tethered to the nuclear surface (Fig. 5E). Therefore, the stoichiometric model of cortical and cytoplasmic pulling forces is promising for centrosome spacing in *C. elegans* embryos and can be applied to other cell types and species.

A similar pulling-force-based mechanism was proposed for the spacing of nuclei in the syncytium embryo of the cricket *Gryllus bimaculatus* (Donoughe et al., 2022), despite the observation of a pushing-based mechanism for similar nuclear spacing in *Drosophila* syncytium embryos (Deshpande et al., 2021). The proposed pulling-based mechanism in crickets supports the generality of the mechanism proposed in the present study for *C. elegans* embryos. However, further studies are necessary to clarify pulling-based mechanisms in crickets. The involvement of dynein and other pulling force generators has not yet been demonstrated in crickets. The current argument against the pushing-based mechanism in crickets is that numerical simulation (Dutta et al., 2019) does not correspond to certain aspects of nuclear migration in crickets (Donoughe et al., 2022). It is possible that kinesin-5, PRC1, or kinesin-4 is required for spacing in crickets. The pulling-based model proposed for crickets (Donoughe et al., 2022) is similar to that used in the present study. Unlike our model, which is independent of the nucleus, the model for the cricket assumed occlusion of the microtubule “cloud” by the nucleus as the primary driving force. In *Drosophila*, the centrosome spacing in the syncytium is independent of the nucleus (de-Carvalho et al., 2022), which may also be the case in crickets. In this scenario, a stoichiometric model of cortical and cytoplasmic pulling forces, which does not require nuclei for centrosome spacing, would be more suitable, even for crickets. Finally, in contrast to the cricket model, in which an occlusion between the microtubule “clouds” was assumed without mechanistic bases, the stoichiometric model of cortical and cytoplasmic pulling forces assumes competition based on the reasonable length distribution of the microtubule (i.e., longer microtubules are rare). In this regard, we believe that the stoichiometric model of cortical and cytoplasmic pulling forces is more widely applicable.

### Myosin-dependent centrosome movement

We observed large movement of centrosomes in dynein (*dhc-1*) knockdown embryos (Fig. 6). Previous studies have focused on some of the earliest phenotypes (defects in the centrosome separation, pronuclear migration, and spindle elongation in the 1-cell stage embryo) of *dhc-1* RNAi or mutant embryos (Gönczy et al., 1999; Schmidt et al., 2005; Kimura and Onami, 2005) and did not focus on the later movements of the centrosomes. The timing of the large movement of the centrosomes in *dhc-1* (RNAi) embryos coincided with that of cytokinesis. This timing suggests the involvement of cytoplasmic flow coupled with cytokinesis (White and Borisy, 1983; Khaliullin et al., 2018). This idea was supported by our RNAi experiment on the non-muscle myosin *nmy-2*, a gene responsible for generating cytoplasmic flow (Shelton et al., 1999). We confirmed that cytoplasmic flow occurred in *dhc-1* (RNAi) cells.

The involvement of the cytoplasmic flow in the movement of the centrosomes (with or without nuclei) has been reported in previous studies. In the 1-cell stage *C. elegans* embryo, soon after symmetry breaking, the centrosomes move along the cortex in *zyg-12* (RNAi) embryos in an *nmy-2*-dependent manner (De Simone et al., 2016). The dependency on *nmy-2* suggests that the driving force for this movement is cytoplasmic flow. However, another interpretation is possible. Because *nmy-2* (RNAi) impaired cortical pulling forces (Redemann et al., 2010), and defective cortical pulling forces impaired movement along the cortex (Fig. 5C and S4, compare control-vs-*gpr-1/2*(RNAi)), the centrosome movement behavior in *zyg-12*; *nmy-2* (RNAi) (De Simone et al., 2016) can be explained by defects in the cortical pulling force.

Another example of centrosome movement by cytoplasmic flow is from the 1-cell stage *C. elegans* embryo, but earlier than symmetry breaking. The cytoplasmic flow, driven by kinesin and microtubules (McNally et al., 2010; Kimura et al., 2017), moves the sperm-derived pronucleus together with the centrosome, affecting the formation of the anterior-posterior axis of the embryo (Kimura and Kimura, 2020). In *Drosophila* syncytium embryos, the movement of nuclei via myosin-dependent cytoplasmic flow is important for nuclear positioning and synchronized cell division (Dassow and Schubiger, 1994; Deneke et al., 2019). Our finding of centrosome movement by cytoplasmic flow may provide insight into how cytoplasmic stream flows into the nucleus and centrosomes in future studies.

## Conclusion

We propose a simple and reasonable model for centrosome spacing that is independent of the nucleus or spindle in *C. elegans* embryos. This mechanism is expected to function in other cell types and organisms in combination with repulsive pushing between centrosomes using antiparallel microtubules. Centrosome spacing according to the stoichiometric model of cortical and cytoplasmic pulling forces may play a role in the rapid placement of centrosomes during development. This may apply to species with asters formed only by short microtubules. The proposed model is based on experiments with enucleated *C. elegans* embryos. An enucleated *C. elegans* embryo is a powerful model for cell and developmental biology, and our experimental setup should be sufficiently powerful to address other biological questions.

## Materials and Methods

### Worm strains and RNAi

The *C. elegans* strains used in this study are listed in Supplementary Table S1. The DE90 strain (*tbg-1*::*GFP*; *GFP*::*histone H2B*; *GFP*::*PH ^PLCδ1^*) was used to obtain embryos with nuclei (*zyg-12, dhc-1* and *dhc-1*; *nmy-2* RNAi experiments). The strains were maintained under standard conditions (Brenner, 1974). Knockdown of *klp-18, zyg-12, gpr-1/2, dhc-1, dyrb-1*, and *nmy-2* was performed using the injection RNAi method as previously described (Kimura and Kimura, 2011). For double- or triple-RNAi experiments, RNA was mixed in a 1:1 or 1:1:1 ratio and injected into the worms. The dsRNA concentrations were 18 or 21 μg/μL for *klp-18*, 15 or 19 μg/μL for *gpr-1/2*, 19 μg/μL for *dhc-1*, and 13 μg/μL for *dyrb-1*. To efficiently obtain the *klp-18* phenotype (enucleated embryos), observations were started ≧24 h after injection. Observations were also conducted at ≧26 h after double knockdown and ≧30 h after triple knockdown. The worms were incubated at 25 °C for ≧16 h before observation (*zyg-12, dhc-1* and *dhc-1*; *nmy-2* RNAi experiments).

### Production of enucleated embryos

Enucleated embryos were produced as follows (Fig. S1). First, 2 young adults of each of the CAL0051, CAL0181, or CAL2741 strains were transferred to a new plate 5 days before the day 1, and preculture was initiated. On day 1, 15 CAL0051 and 10 CAL0181 or CAL2741 young adults were separately moved onto new 6 cm (diameter) NGM plates with the OP50 *E. coli,* and the plates were cultured at 16 °C, a non-restrictive temperature. After 24 h, on day 2, the worms were removed from the plate, and only embryos that had been laid in the last 24 h remained on the plate. Cultures were maintained at 16 °C. On day 3, 24 h after the procedures on day 2, the plates were transferred to 25 °C, which is a restrictive temperature. On day 4, 24 h after the procedures on day 3, 25 CAL0181 or CAL2741 L4 or young adults were selected (hermaphrodites with vulva) and injected with *klp-18* dsRNA. After injection, culturing on the NGM plate was continued, with 5 times the number of CAL0051 males added (e.g. 25 hermaphrodites and 125 males). A 3.5-cm NGM plate was used for mating. Finally, on day 5, 24 h or more after the injection, the worms were dissected, and the embryos were observed under a fluorescence microscope.

### Microscopic observation

The localization of the fluorescent proteins was observed using a spinning-disk confocal microscope (CSU-MP; Yokogawa Electric, Tokyo, Japan) (Otomo et al., 2015; Kamada et al., 2022) equipped with an and an EM-CCD camera (iXon; Andor, Belfast, UK) mounted on an IX71 microscope (Olympus, Tokyo, Japan) and controlled using NIS-elements software (Nikon, Tokyo, Japan). Details of the system and the examination of phototoxicity will be published elsewhere (in preparation). Dissected worm embryos were attached to a poly-L-lysine-coated cover glass, mounted under the microscope and observed using a 40× objective lens with 2× intermediate magnification.

To analyze the centrosome distance, two-photon excitation with a 920 nm laser (ALCOR920-2, Spark Lasers, Martillac, France) was used with 96-ms exposure. For Fig. 1, 2, 3, and related supplemental figures and movies, 61 or 71 images were taken at 0.5 μm intervals on the z-axis. Time-lapse images were collected at 1 min intervals for less than 2 h. For Fig. 6, 41 images were taken at 0.5 μm intervals on the z-axis. Time-lapse images were collected at 10 s intervals during the 1-cell stage.

To analyze the distribution of microtubules and centrosomes (Fig. 4 and S3), the dissected worm embryos were attached to a 2% agarose-coated cover glass and mounted on the microscope. Single focal plane was captured using a single photon excitation with a 488 nm laser with 804-ms exposure. The focal plane was manually adjusted to the target aster during this interval. Time-lapse images were collected at ∼1-min-interval operated manually for 30 min.

Under these microscopic conditions, *C. elegans* embryos were confirmed to hatch. The captured images were analyzed using the ImageJ/Fjii or Imaris software (Oxford instruments).

### Measurement of centrosome distance

The distance between the centrosomes was quantified using the Imaris 3D analysis software. The centrosome signals were tracked manually using the spot-tracking mode. The centroid coordinates of the centrosome signals were calculated by the software. From the calculated coordinates, the distance between centrosomes was calculated. For the wild-type, *zyg-12* (RNAi), and enucleated embryos (Fig. 2C, D, and 3B), the centrosomes (Fig. 2A), which split into two in the 2 cell stage, were tracked until the signal became undetectable or until the next duplication occurred. For Fig. 6, centrosome signals during the 1-cell stage were tracked until they became undetectable.

### Analysis of microtubule distribution

For β-tubulin::GFP images (Fig. 4A), the center coordinates of the centrosomes were quantified using the SpotTracker plugin in ImageJ/FIJI (Sage et al., 2005) (http://bigwww.epfl.ch/sage/soft/spottracker/). The fluorescence intensity of soluble β-tubulin was defined as the peak intensity of the cytoplasmic signal. The fluorescence intensity of polymerized β-tubulin (i.e. microtubules) was defined as the captured intensity subtracted by the soluble β-tubulin intensity. The mean and S.E.M. of the subtracted intensity were calculated for the ring-shaped region for every 4-pixel thickness. The mean intensity was multiplied by the average circumference of the ring. The summed intensity of the ring regions, *S*(*R*), should be proportional to the number of microtubules reaching the rings and was plotted against the average radius of the ring, *R* (Fig. 4B). The plot was fitted with a combination of two functions: *S*(*R*) = *a*×*R* (for *R* < *R*_0_) and *S*(*R*) = *a*×*R*_0_×EXP[-{(*R-R*_0_)/*ξ*}^*P*] (for *R* ≧ *R*_0_), where *a, R*_0_, *ξ* and *P* are the fitting parameters. The fitting was conducted with a maximum likelihood method (Yesbolatova et al., 2022), assuming that the error was normally distributed with its mean summed-intensity and variance as the square of the standard error of the mean (S.E.M.) multiplied by the circumference length, using the solver function of Microsoft Excel (Microsoft Corporation). The function *S*(*R*) = *a*×*R*_0_×EXP[-{(*R-R*_0_)/*ξ*}^*P*] represents the Weibull distribution. The Weibull distribution is used to model the distribution of the lifespan, whose death rate is proportional to the power of time. We confirmed that the simulated microtubule length distribution of constant growth/shrinkage velocity and catastrophe/rescue frequency fit well with the Weibull distribution. Therefore, we determined the length (*l*) distribution of the microtubule as *S*(*l*) = *a*×*l*_0_×EXP[-{(*l-l*_0_)/*ξ*}^*P*] (for *l* ≧ *l*_0_, where *l*_0_ is the radius of the centrosome).

To calculate the average distribution of the microtubule length at every time point and sample, we first fitted the microtubule length distribution at each time point for each sample using the Weibull distribution. After fitting, the average value of the fitted distribution was calculated to obtain the average distribution. This average distribution was further fitted to the Weibull distribution to obtain the function parameters. The microtubule length distribution in our simulation was obtained using the Weibull distribution.

### Statistical analysis

The distances between the centrosomes were statistically compared using the Wilcoxon rank sum test, which was performed for the two experimental groups of interest. Calculations were performed using the “ranksum” function of MATLAB software (The Mathworks).

### Numerical simulation of the stoichiometric model of cortical and cytoplasmic pulling forces

The settings of our previous simulation (Kondo and Kimura, 2019) were modified to model the embryo as a 3D ellipsoid with the long axis of 46.0 μm and the two short axes of 27.6 μm based on the size of a representative enucleated (control) embryo. For the simulation of gene knockdown conditions, we modified the sizes based on the sizes of each condition (Table S3). As in the previous simulation, we distributed “force generation points” throughout the cytoplasm and the cortex (3 μm thick layer) at the vertices of a simple cubic lattice with 1 μm intervals. When a force generation point was associated with a microtubule elongating from the centrosome, it pulled the centrosome with a defined force (Table S2).

The probability of the point attaching a microtubule from the *i*-th centrosome was defined by the distance between the point and the centrosomes, as assumed in the stoichiometric model of cortical pulling forces proposed by Farhadifar et al. (2020): 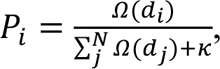, where *Ω*(*d*_i_) is the rate microtubules from *i*-th centrosome contact a force-generator at a distance of *d*_i_, and *κ* is the rate a force generator detach from a microtubule. There were three notable differences between the original stoichiometric model (Farhadifar et al., 2020) and the stoichiometric model of the cortical and cytoplasmic pulling forces in this study. First, the model was extended to simulate the behavior of more than three centrosomes. Second, the force generators pull microtubules in the cortex and the cytoplasm, based on our experimental results that simultaneous knockdown of *gpr-1/2* and *dyrb-1* but not either, is required for the spacing defect comparable to *dhc-1* (RNAi) (Fig. 3). Third, we used the distribution of microtubule lengths based on our own experimental measurements of signal intensity, reflecting the microtubule distance, *d*, from the center of the centrosome: *S*(*d*) = *a*×*d*_0_×EXP[-{(*d-l*_0_)/*ξ*}^*P*] (for *d* ≧ *l*_0_), where *l*_0_ is the radius of the centrosome (Fig. 4 and Table S2). Finally, we defined *Ω*(*d*) as *Ω*(*d*) = (*γ*/4)(*r*/*d*)^2^×EXP[-{(*d-l*_0_)/*ξ*}^*P*] (for *d* ≧ *l*_0_), where *γ* is the rate of microtubule nucleation at the centrosome and *r* is the force-generator capture radius. For the case where the force generator is located inside the centrosome (*d* < *l*_0_), we assumed that all the nucleated microtubules reach the distance, and thus defined *Ω*(*d*) as *Ω*(*d*) = (*γ*/4)(*r*/*d*)^2^ (for *d* < *l*_0_).

Once we calculated the probability *P*_i_, for each force generator to pull the *i*-th centrosome, the force that pulls the *i*-th centrosome was calculated as *f*_0_*P*_i_, where *f*_0_ is the force generated by each force generator (Farhadifar et al., 2020). In this study, we define *f*_0_cort_ and *f*_0_cyto_ as the forces generated by the cortical and cytoplasmic force generators, respectively. The total force vector acting on each centrosome was calculated by summing all the force vectors of the force generators and pulling the centrosome toward the direction of each force generator. After summing the forces acting on each centrosome, the velocity of the movement was calculated as 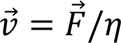, where 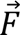, *η*, and 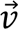 are the force, drag coefficient, and velocity vector, respectively. The positions of the centrosomes in the next step were calculated as 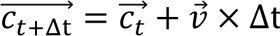, where 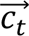 and 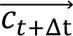 are the position vectors of the centrosomes at times *t* and *t*+Δ*t*, respectively, and Δ*t* is the time interval. This calculation was repeated for the defined steps starting from the initial positions of the centrosomes.

In the case where the centrosomes were tethered to the surface of the nucleus, we added an additional process after each step to apply an elastic force, 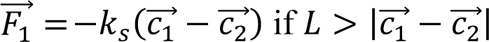 to maintain the distance between the centrosomes at *L* or shorter. Here, 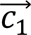 and 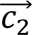 are the position vectors of the centrosome, force applied, and other centrosomes, respectively. *k_s_* is the elastic constant and *L* is the diameter of the nucleus (8 μm).

The simulation was coded using MATLAB, and the codes are available upon request.

### The parameter values of the numerical simulation

The parameter values used for the numerical simulations are summarized in Supplemental Tables S2 and S3. We followed the embryo geometry based on the experimental measurements and the simulation setup in our previous study (Kondo and Kimura, 2019). The parameters for force generation are basically the same as those of Farhadifar et al. (2020). In our setup, the number of cortical force generators (*N*_cort_) was 12,408 for the control condition. According to Farhadifar et al. (2020), the pulling force produced by a force generator (*f*_0_cort_) was 0.08 pN, the force-generator capture radius (*r*) was 0.1 μm, and the microtubule-force-generator detachment rate (*κ*) was 4.4×10^-4^/s.

To determine the cortical pulling force, cytoplasmic pulling force, and force reduction by RNAi, we compared the simulated results of the distance between sister centrosomes under different conditions (control, reduced cortical pulling force, and reduced cytoplasmic pulling force) with the corresponding experimental results (control, *gpr-1/2, dyrb-1*, as shown in Fig. 3B and S6). First, we searched for appropriate values for the force produced by a cortical force generator (*f*_0_cort_) and a cytoplasmic force generator (*f*_0_cyto_) that reproduced the maximum rate of increase in distance for *dyrb-1* (RNAi)-enucleated embryos (i.e., defective cytoplasmic forces) and *gpr-1/2* (RNAi)-enucleated embryos (i.e., defective cortical forces), respectively. (The number of cytoplasmic force generators (*N*_cyto_) in our setup was 18,389 for the control condition). The average values of the optimized force parameters for the three pairs of representative embryos were 0.034 pN for *f*_0_cort_ and 0.014 pN for *f*_0_cyto_. Using these parameter values, we simulated centrosome movement in control, *dyrb-1* (RNAi), *gpr-1/2* (RNAi), and *dhc-1* (RNAi) enucleated embryos. To mimic the low-level spacing in *dhc-1* (RNAi)-enucleated embryos, we assumed that RNAi treatments reduced the force (*f*_0_cort_ and *f*_0_cyto_) to 5% but not to 0%.

## Acknowledgments

We thank Ms. Tomoko Ozawa for preparing the experimental tools and Ms. Yoko Kimura for establishing the worm strains used in this study. We thank Drs. Hitoshi Sawa, Hironori Niki, Masato Kanemaki, Yuta Shimamoto, and Hirokazu Tanimoto, the members of the Cell Architecture Laboratory and Physics and Cell Biology Laboratory of National Institute of Genetics, for their fruitful discussions.

## Author Contributions

K.F, T.K, and A.K conceived the study. K.F, T.K, and A.K designed the experiments. K.F and T.K performed the experiments and analyzed the data. A.K wrote the computational codes. K.F. ran the codes and analyzed the data. K.F and A.K wrote the manuscript. K.F, T.K, and A.K reviewed and edited the manuscript.

## Supplemental Tables

**Supplemental Table S1.** Strains used in this study.

| Strain | Genotype | Cultivation temperature |
| --- | --- | --- |
| CAL0181 | *fem-1(hc17ts) IV* (Temperature sensitive).<br>*ruIs32* [*pie-1p::GFP::histone H2B* + *unc-119(+)*]*III*.<br>*ddIs6* [*tbg-1::GFP* + *unc-119(+)*] *V*.<br>*ltIs38* [*pie-1p::GFP::PH(PLC1 δ 1)* + *unc-119(+)*]. | 16 °C |
| CAL0051 | *emb-27(g48ts) II* (Temperature sensitive).<br>*him-5(e1490) V*. | 16 °C |
| CAL2741 | *fem-1(hc17ts) IV* (Temperature sensitive).<br>*ruIs32* [*pie-1p::GFP::histone H2B* + *unc-119(+)*]*III*.<br>*ojIs1* [*pie-1p::GFP::tbb-2* + *unc-119(+)*]. | 16 °C |
| DE90 | *unc-119(ed3 or e2498)*<br>*oxIs318* [*spe-11p::mCherry::histone* + *unc-119(+)*] *II*.<br>*ruIs32* [*pie-1p::GFP::histone H2B* + *unc-119(+)*] *III*.<br>*ddIs6* [*tbg-1::GFP* + *unc-119(+)*] *V*.<br>*dnIs17* [*pie-1p::GFP::PH(hPLCIIIδ1)* + *unc-119(+)*]. | 16 or 22 °C |

**Supplemental Table S2:**
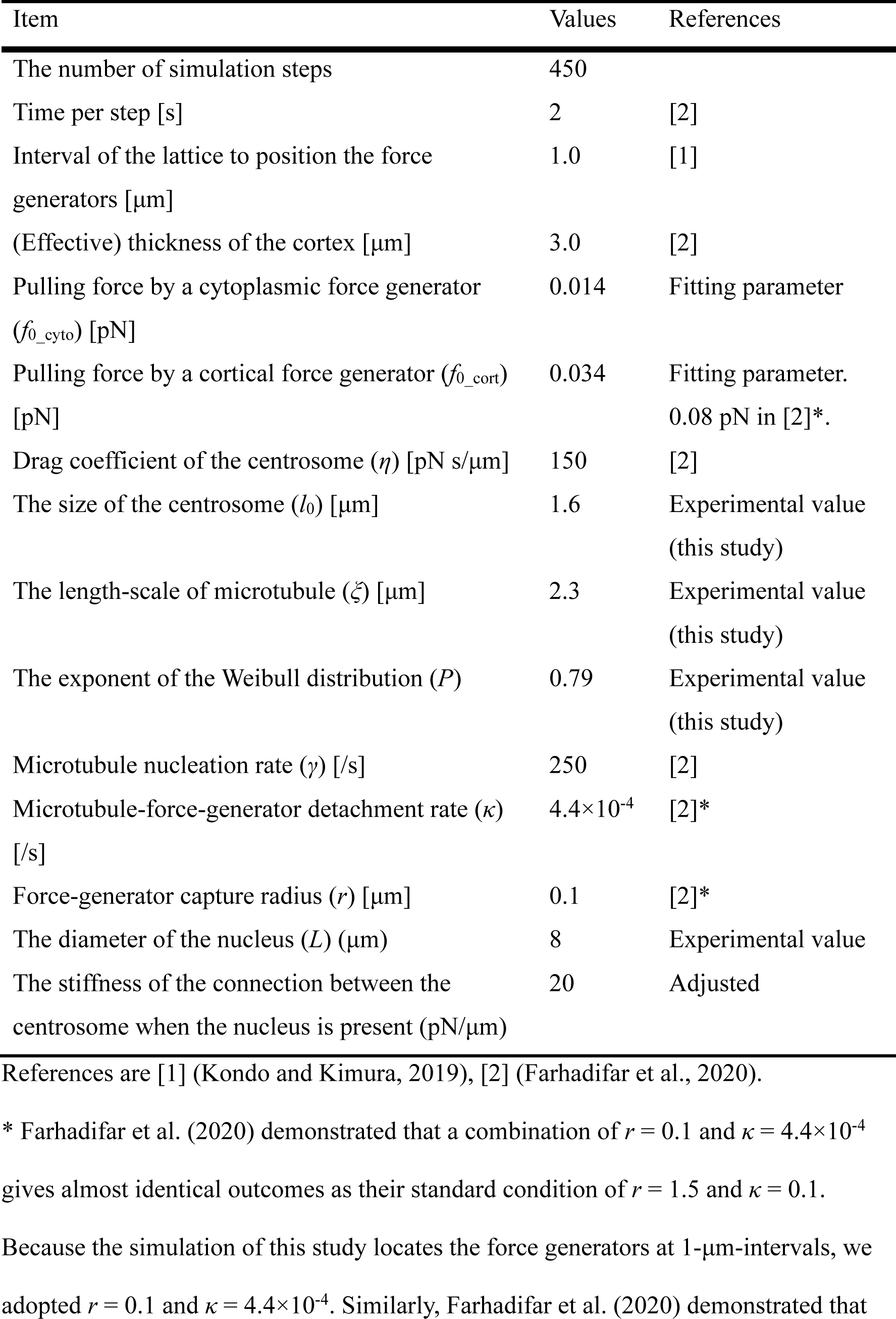
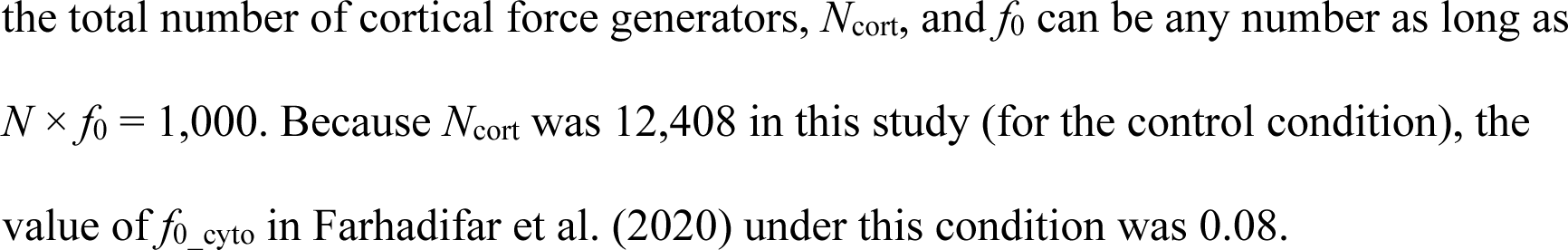
parameters for the simulations.

| Item | Values | References |
| --- | --- | --- |
| The number of simulation steps | 450 |  |
| Time per step [s] | 2 | [2] |
| Interval of the lattice to position the force generators [$\mu\text{m}$] | 1.0 | [1] |
| (Effective) thickness of the cortex [$\mu\text{m}$] | 3.0 | [2] |
| Pulling force by a cytoplasmic force generator ($f_{0\_cyto}$) [pN] | 0.014 | Fitting parameter |
| Pulling force by a cortical force generator ($f_{0\_cort}$) [pN] | 0.034 | Fitting parameter.<br>0.08 pN in [2]*. |
| Drag coefficient of the centrosome ($\eta$) [pN s/$\mu\text{m}$] | 150 | [2] |
| The size of the centrosome ($l_0$) [$\mu\text{m}$] | 1.6 | Experimental value<br>(this study) |
| The length-scale of microtubule ($\xi$) [$\mu\text{m}$] | 2.3 | Experimental value<br>(this study) |
| The exponent of the Weibull distribution ($P$) | 0.79 | Experimental value<br>(this study) |
| Microtubule nucleation rate ($\gamma$) [/s] | 250 | [2] |
| Microtubule-force-generator detachment rate ($\kappa$) [/s] | $4.4 \times 10^{-4}$ | [2]* |
| Force-generator capture radius ($r$) [$\mu\text{m}$] | 0.1 | [2]* |
| The diameter of the nucleus ($L$) ($\mu\text{m}$) | 8 | Experimental value |
| The stiffness of the connection between the centrosome when the nucleus is present (pN/$\mu\text{m}$) | 20 | Adjusted |

References are [1] (Kondo and Kimura, 2019), [2] (Farhadifar et al., 2020).
\* Farhadifar et al. (2020) demonstrated that a combination of $r = 0.1$ and $\kappa = 4.4 \times 10^{-4}$ gives almost identical outcomes as their standard condition of $r = 1.5$ and $\kappa = 0.1$ .
Because the simulation of this study locates the force generators at 1- $\mu\text{m}$ -intervals, we adopted $r = 0.1$ and $\kappa = 4.4 \times 10^{-4}$ . Similarly, Farhadifar et al. (2020) demonstrated that
the total number of cortical force generators, $N_{\text{cort}}$ , and $f_0$ can be any number as long as $N \times f_0 = 1,000$ . Because $N_{\text{cort}}$ was 12,408 in this study (for the control condition), the value of $f_{0\_cyto}$ in Farhadifar et al. (2020) under this condition was 0.08.

**Supplemental Table S3:** The condition-dependent parameters of the simulations.

| Condition | Size of the embryo:<br>long axis × short axis [$\mu\text{m}$]<br>number of force generators* | Initial coordinates of<br>the centrosomes<br>[$\mu\text{m}$] |
| --- | --- | --- |
| control embryo | $46.0 \times 27.6$ | (17.4, 2.8, 0.8) |
| (cortexF $\times 1$, cytoF $\times 1$) | $N_{\text{cort}} = 12,408$ | (18.1, -1.3, -0.3) |
| | $N_{\text{cyto}} = 18,389$ | (-7.0, -1.1, 0.1) |
|  |  | (-7.2, 1.6, 0.9) |
| *gpr-1/2* (RNAi) embryo | $50.6 \times 28.6$ | (19.7, 1.0, 5.3) |
| (cortexF $\times 1/20$, cytoF $\times 1$) | $N_{\text{cort}} = 13,974$ | (20.1, 1.1, 6.7) |
| | $N_{\text{cyto}} = 21,617$ | (14.2, -3.9, -6.5) |
|  |  | (13.0, -3.3, -7.0) |
|  |  | (-6.3, -0.3, -0.8) |
|  |  | (-7.2, -0.0, -1.4) |
| *dyrb-1* (RNAi) embryo | $49.2 \times 28.0$ | (20.6, -1.7, 2.0) |
| (cortexF $\times 1$, cytoF $\times 1/20$) | $N_{\text{cort}} = 13,096$ | (20.5, -2.4, 0.5) |
| | $N_{\text{cyto}} = 20,171$ | (1.2, -7.5, 3.9) |
|  |  | (-0.1, -7.6, 4.1) |
|  |  | (-10.8, 4.4, 2.1) |
|  |  | (-11.1, 5.9, 3.6) |
| *dhc-1* (RNAi) embryo | $49.2 \times 27.4$ | (17.1, -7.7, 9.2) |
| (cortexF $\times 1/20$, cytoF $\times 1/20$) | $N_{\text{cort}} = 13,092$ | (17.9, -7.5, 9.2) |
| | $N_{\text{cyto}} = 19,321$ | (17.2, -12.1, -3.8) |
|  |  | (18.0, -11.4, -4.3) |
|  |  | (18.0, 9.5, -4.1) |
|  |  | (17.6, 9.2, -5.1) |

\* $N_{\text{cort}}$ is the number of cortical force generators, and $N_{\text{cyto}}$ is the number of cytoplasmic force generators. The both depend on the cell size.

## Supplemental Figures

**Supplemental Fig S1:**
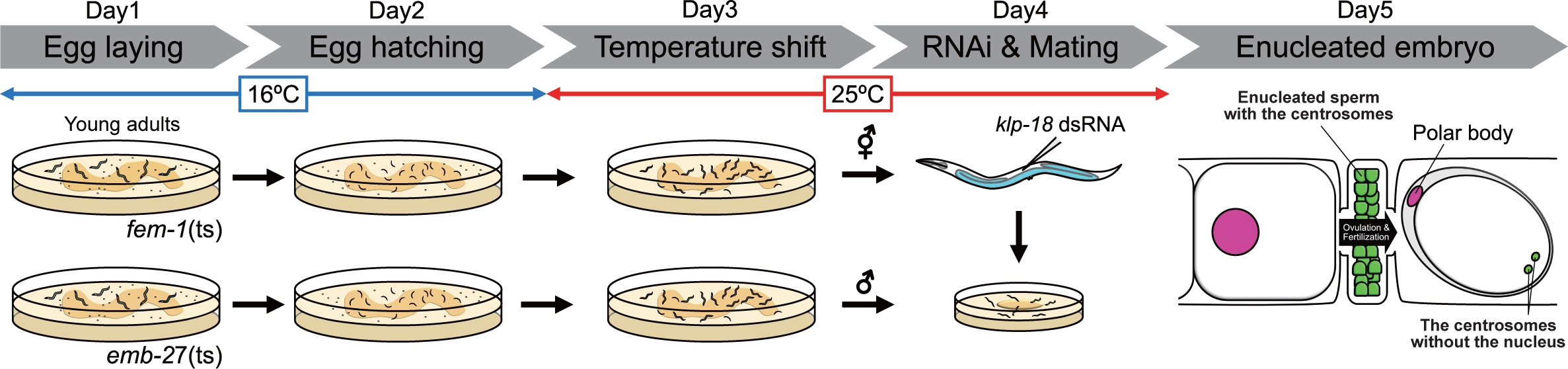
The procedure of enucleation of the *C. elegans* embryo. Related to Fig. 1. Schematic drawing of the enucleation procedure. For the details, see Materials and Methods. Day 1: Young adults were moved onto fresh culture plates to lay eggs. Day 2: The adults were removed from the plate. Day 3: The plates were transferred from 16℃ to 25℃. Day 4: The hermaphrodites were injected with *klp-18* dsRNA. After injection, the hermaphrodites were cultured on a smaller plate containing the males. Day 5: The hermaphrodites were dissected and observed under the fluorescence microscope.

**Supplemental Fig. S2:**
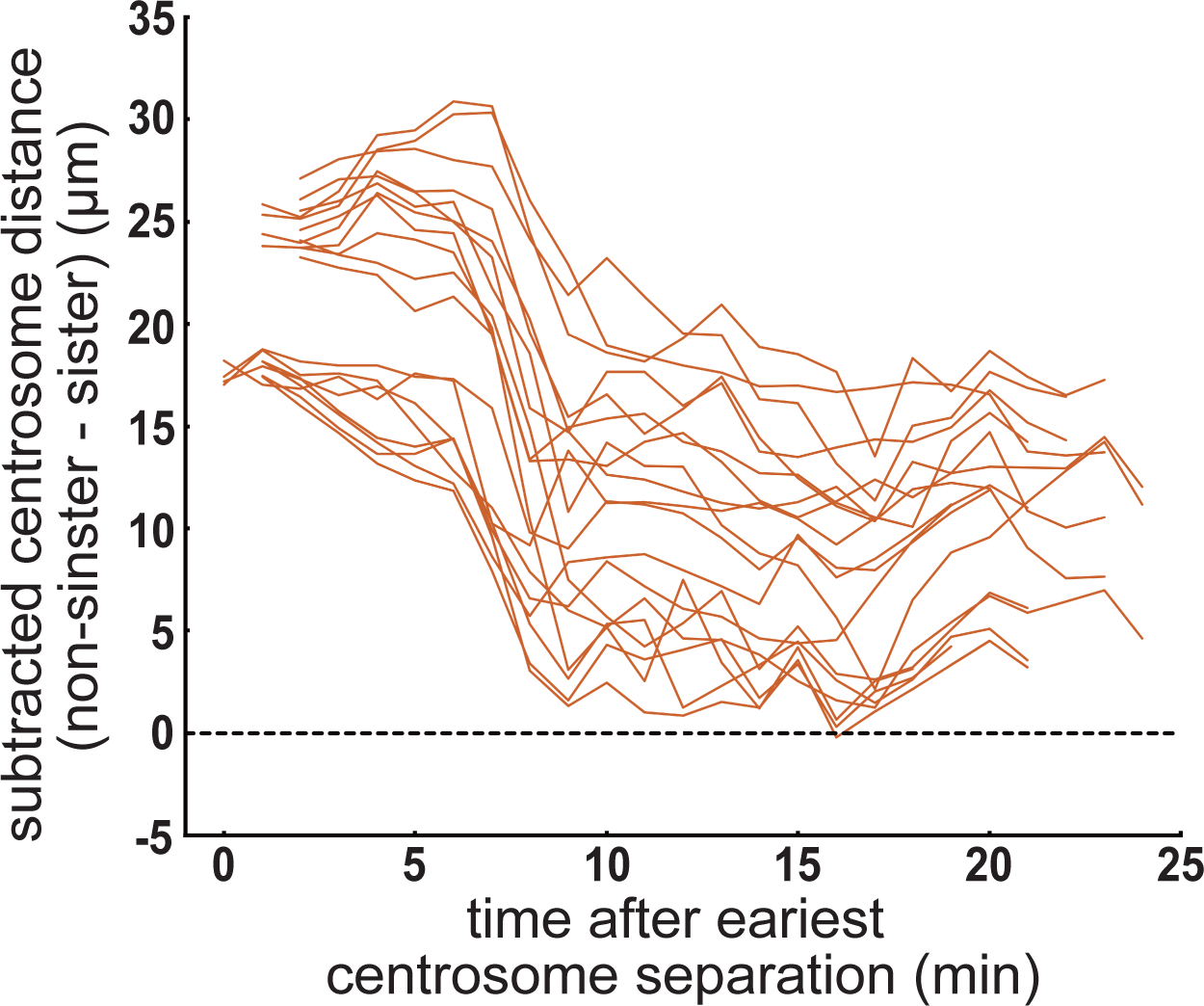
Characterization of centrosome dynamics during the second cell cycle in enucleated embryos. Related to Fig. 2. Distance transition between non-sister centrosomes. The minimum sister centrosome distance was subtracted from the distance between non-sister-pairs of centrosomes at each time point in the enucleated embryo. Individual samples are indicated with red lines (20 pairs from 5 embryos). The black dotted line indicates the subtracted distance = 0. The distances between non-sister pairs were rarely shorter than those between sister pairs, indicating that a similar spacing mechanism was applied for both sister and non-sister centrosome pairs.

**Supplemental Fig. S3:**
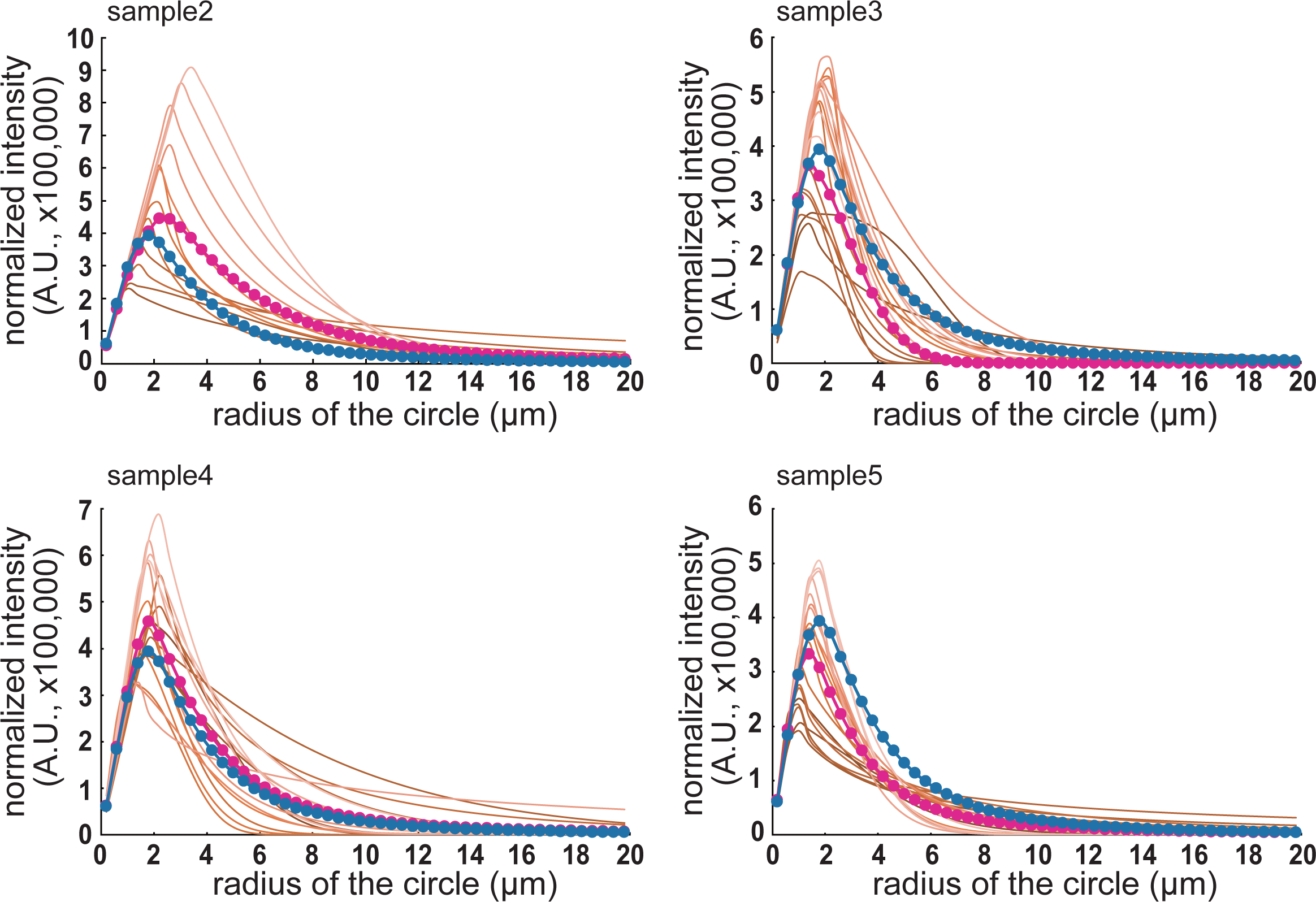
Distribution of the microtubule length in enucleated embryos. Related to Fig. 4. The fitting results of the β-tubulin signals in 4 enucleated embryo, other than the one shown in Fig. 4C (“sample 1”). The fitted curves at each time point are indicated by the brown lines. Darker colors indicate earlier time points. Lighter colors indicate later time points. The average fitting curve is shown with magenta dots and lines. The average fitting curve for the 5 embryos is indicated by blue dots and lines. We concluded that the estimated distribution of microtubule lengths did not change dramatically among the different samples.

**Supplemental Fig. S4:**
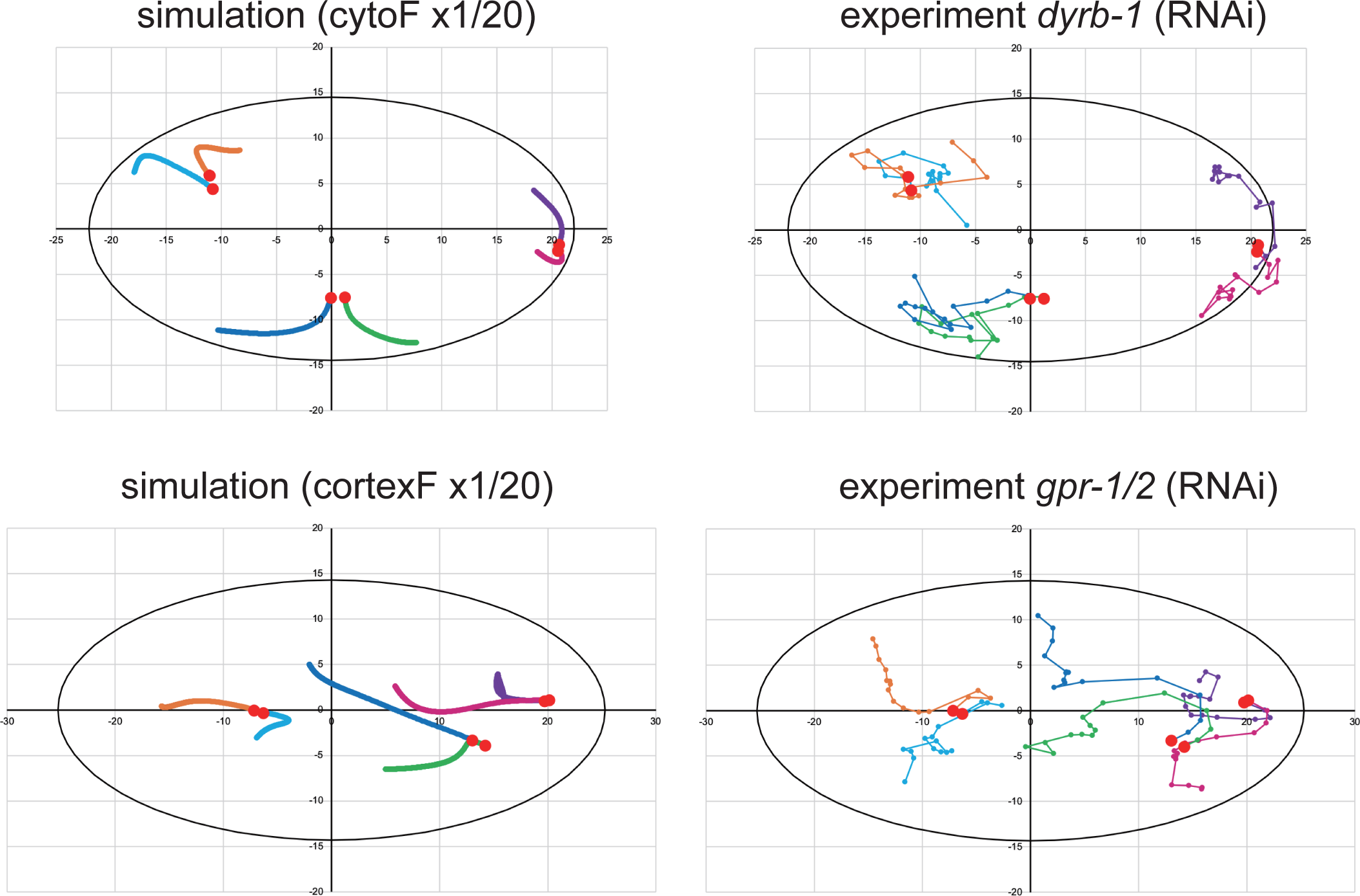
Trajectories of the centrosomes in the simulation and experiment. Related to Fig. 5. Trajectories of the centrosomes in a representative enucleated embryo (right) and a simulation with the same initial positions as the six centrosomes (left). Red circles indicate the initial positions of the centrosomes. Trajectories of the same color indicate the same initial positions. The trajectories of the centrosomes inside the cell were similar in the simulations and experiments.

**Supplemental Fig. S5:**
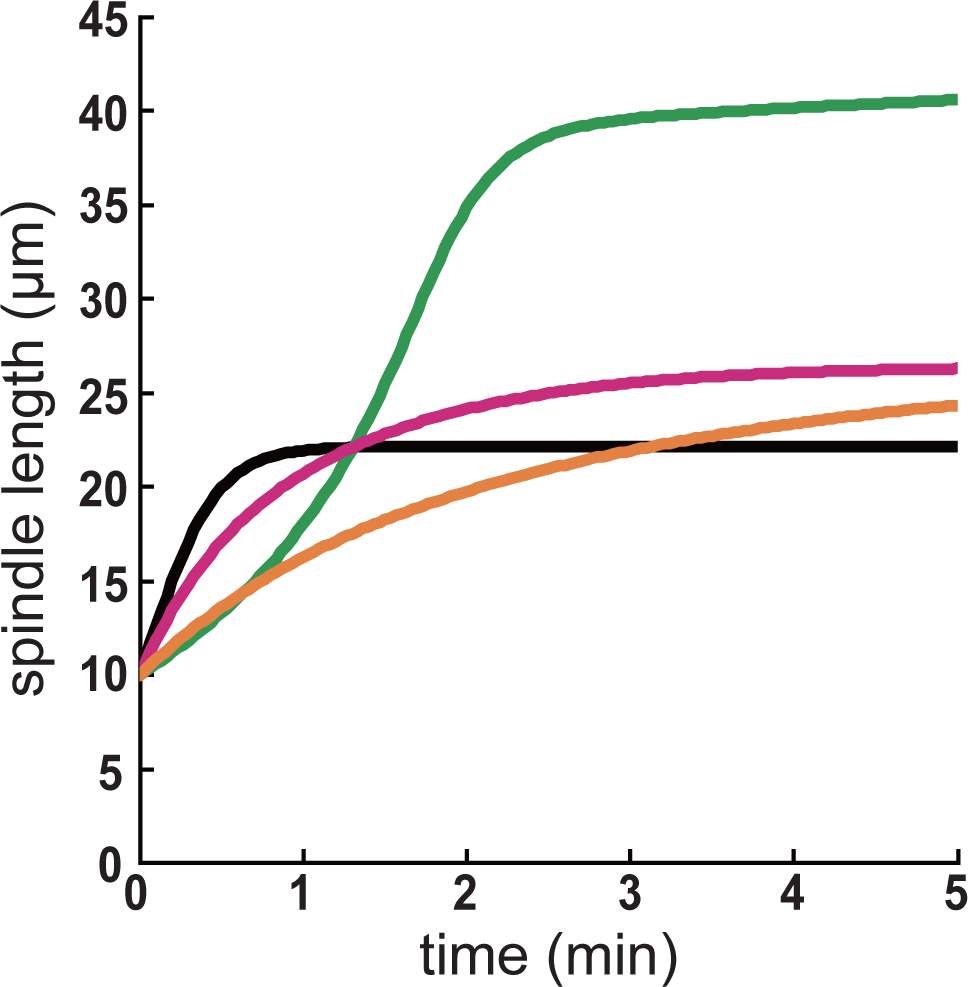
Stoichiometric model of cortical and cytoplasmic pulling forces reproduces spindle elongation. Related to Fig. 5. Simulated results for the spindle length. The line indicates the simulated spindle length. Black: Simulation only with cortical pulling forces (*f_0_cort_* = 0.08), without cytoplasmic pulling forces (*f_0_cyto_* = 0), and with long microtubules (exponential decay with a characteristic length of 20 μm) as assumed in Farhadifar et al. (2020). Green: Simulation as in Black except using the experimentally obtained microtubule distribution (Fig. 4D and Table S2). Magenta: Simulation as in Green except adding cytoplasmic pulling forces (*f_0_cyto_* = 0.033). Orange: Simulation as in Magenta except using the force parameters as same as in our other simulation (*f_0_cort_* = 0.034 and *f_0_cyto_* = 0.014, Fig. 5, S4, and Table S2).

**Supplemental Fig. S6:**
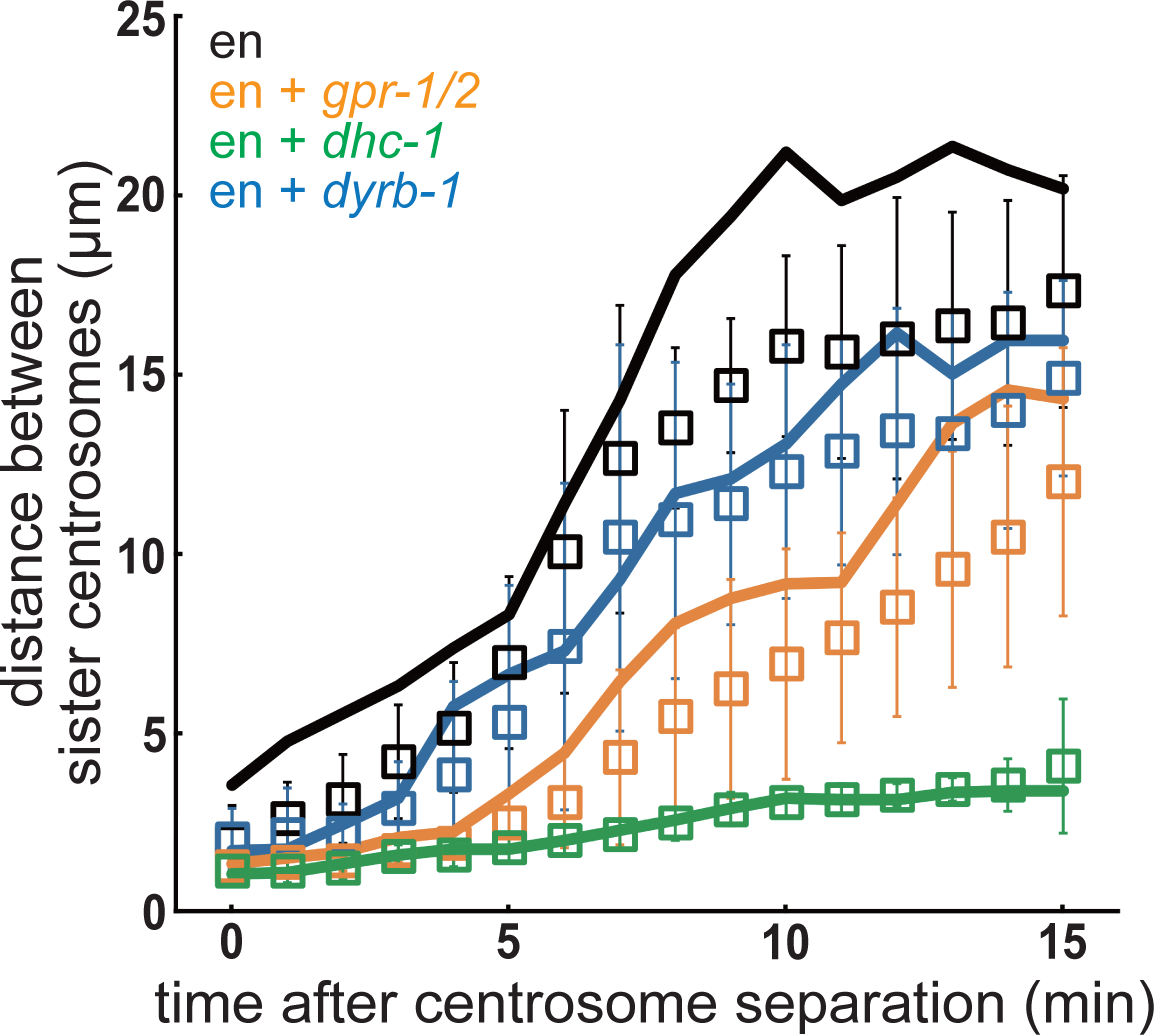
Mean distance between sister centrosomes in the representative enucleated embryos. Related to Fig. 5. The mean distance between sister centrosomes in a representative embryo from each condition is shown with solid lines. The mean and S.D. of all embryos, which are identical to the results shown in Fig. 3B, are shown as squares and error bars, respectively. Black, enucleated embryos. Orange, *gpr-1/2* (RNAi) enucleated embryos. Green, *dhc-1* (RNAi) enucleated embryos. Blue, *dyrb-1* (RNAi) enucleated embryos.

## Supplemental Movies

**Supplemental Movie S1: Centrosome movement and cell division in control *C. elegans* embryos.**

Time-lapse movie corresponding to Fig. 1A. Time-lapse movie of *C. elegans* embryos expressing *GFP*::*histone H2B, tbg-1*::*GFP, GFP*::*PH^PLC1δ1^*. In the first 5 frames, the yellow arrows indicate the centrosomes, yellow arrowheads indicate the pronuclei, and yellow circles indicate the polar bodies. The movie of 2-hour imaging is shown. z-maximum projections. Time is indicated in min. Time 0 was when the imaging started. The time interval between measurements was 1 min. Scale bar, 10 μm.

**Supplemental Movie S2: Centrosome movement and cell division in *emb-27*(*g48ts*) mutant *C. elegans* embryos.**

Time-lapse movie corresponding to Fig. 1B. Imaging condition was same as in Movie S1.

**Supplemental Movie S3: Centrosome movement and cell division in *klp-18* (RNAi) *C. elegans* embryos.**

Time-lapse movie corresponding to Fig. 1C. Imaging condition was same as in Movie S1.

**Supplemental Movie S4: Centrosome movement and cell division in *emb-27*(*g48ts*) mutant and *klp-18* (RNAi) *C. elegans* embryos (enucleated embryos).**

Time-lapse movie corresponding to Fig. 1D. Imaging condition was same as in Movie S1.

**Supplemental Movie S5: Centrosome movement in control *C. elegans* embryos during 2-cell stage.**

Time-lapse movie corresponding to Fig. 2B (control). The 2-cell stage is shown. In the first 5 frames, yellow arrows indicate representative sister centrosomes. Time 0 was defined as the time at which representative sister centrosomes were detected. Otherwise, imaging condition was same as in Movie S1.

**Supplemental Movie S6: Centrosome movement in *zyg-12* (RNAi) *C. elegans* embryos during 2-cell stage.**

Time-lapse movie corresponding to Fig. 2B (*zyg-12* (RNAi)). Imaging condition was same as in Movie S5.

**Supplemental Movie S7: Centrosome movement in enucleated *C. elegans* embryos during 2-cell stage.**

Time-lapse movie corresponding to Fig. 2B and Fig. 3A (enucleated embryo). Imaging condition was same as in Movie S5.

**Supplemental Movie S8: Centrosome movement in *gpr-1/2* (RNAi) in enucleated *C. elegans* embryos during 2-cell stage.**

Time-lapse movie corresponding to Fig. 3A (*gpr-1/2* (RNAi) enucleated embryo). For this individual, we did not detect the signal of *GFP*::*PH^PLC1δ1^*. Otherwise, imaging condition was same as in Movie S5.

**Supplemental Movie S9: Centrosome movement in *dhc-1* (RNAi) in enucleated *C. elegans* embryos during 2-cell stage.**

Time-lapse movie corresponding to Fig. 3A (*dhc-1* (RNAi) enucleated embryo). Imaging condition was same as in Movie S5.

**Supplemental Movie S10: Centrosome movement in *gpr-1/2*;*dyrb-1* (RNAi) in enucleated *C. elegans* embryos during 2-cell stage.**

Time-lapse movie corresponding to Fig. 3A (*dyrb-1*; *gpr-1/2* (RNAi) enucleated embryo). For this individual, we did not detect the signal of *GFP*::*PH^PLC1δ1^*. Otherwise, imaging condition was same as in Movie S5.

**Supplemental Movie S11: Centrosome movement in *dyrb-1* (RNAi) in enucleated *C. elegans* embryos during 2-cell stage.**

Time-lapse movie corresponding to Fig. 3A (*dyrb-1* (RNAi) enucleated embryo). Imaging condition was same as in Movie S5.

**Supplemental Movie S12: Centrosome movement in *dhc-1* (RNAi) *C. elegans* embryos during 1-cell stage.**

Time-lapse movie corresponding to Fig. 5A (*dhc-1* (RNAi) embryo). The *C. elegans* embryo contains the nuclei and expressing *GFP*::*histone H2B, tbg-1*::*GFP, GFP*::*PH ^hPLCIIIδ1^*. In the first 10 frames, yellow arrows indicate representative sister centrosomes. 1-cell stage imaging movie is shown. z-maximum projections. Time 0 was when the imaging started. The time interval between measurements was 10 s. Scale bar, 10 μm.

**Supplemental Movie S13: Centrosome movement in *dhc-1*;*nmy-2* (RNAi) *C. elegans* embryos during 1-cell stage.**

Time-lapse movie corresponding to Fig. 5A (*nmy-2;dhc-1* (RNAi) embryo). Imaging condition was same as in Movie S12.

